# Broad variation in rates of polyploidy and dysploidy across flowering plants is correlated with lineage diversification

**DOI:** 10.1101/2021.03.30.436382

**Authors:** Shing H. Zhan, Sarah P. Otto, Michael S. Barker

## Abstract

Changes in chromosome number are considered an important driver of diversification in angiosperms. Single chromosome number changes caused by dysploidy may produce strong reproductive barriers leading to speciation. Polyploidy, or whole genome duplication, yields new species that are often reproductively isolated from progenitors and may exhibit novel morphology or ecology that may further facilitate diversification. Here, we examined the rates of polyploidy, dysploidy, and diversification across the angiosperms. Our analyses of nearly 30,000 taxa representing 46 orders and 147 families found that rates of polyploidy and dysploidy differed by two to three orders of magnitude. The rates of polyploidy and dysploidy were positively correlated with diversification rates, but relative importance analyses indicated that variation in polyploidy was better correlated with diversification rates than dysploidy. Our results provide an overview of angiosperm chromosomal evolution and a roadmap for future research on the complex relationships among polyploidy, dysploidy, and diversification.

## INTRODUCTION

Chromosome number is a dynamic feature of plant evolution, especially among the flowering plants, or angiosperms ^1^. Somatic chromosome counts range from 2*n* = 4 in some angiosperms (e.g., *Colpodium versicolor*) ^2^ to 2*n* = 1,262 in a fern, *Ophioglossum reticulatum* ^3^. Changes in chromosome number occur via polyploidy (whole genome duplication) or dysploidy (single chromosome karyotypic changes). These mechanisms of chromosomal evolution have contributed to the rich diversity of angiosperm genome organization ^4^.

Polyploidy is proposed to underlie the evolution of key innovations ^5^, alter physiological and ecological adaptations ^6–11^, and influence lineage diversification in the angiosperms ^12–14^. Indeed, polyploidy is pervasive throughout the history of the angiosperms ^15^. Polyploidy has been correlated with some upticks in lineage diversification in the long term ^13, 14, 16^ but tends to be associated with lower diversification rates in the short term ^17, 18^. Given the potential deleterious effects of polyploidy (e.g., reduced hybrid fertility ^19^ and increased masking of alleles from selection ^20–22)^, it is still debated whether polyploidy is an evolutionary “dead-end” (*sensu* Stebbins ^23^ and Wagner ^24^ or *sensu* Mayrose et al. ^17, 25^ and Arrigo and Barker ^17, 25^) or brings an occasional boon to angiosperm lineages, perhaps during moments of environmental upheaval ^7, 9^. Recent analyses indicate that polyploidy has a complex relationship with lineage diversification that is mediated by its interactions with other biological factors, such as selfing rate ^16, 26^.

Compared to polyploidy, dysploidy has enjoyed far less attention. Dysploidy is a change in chromosome number (up or down, typically by one chromosome), caused by structural rearrangements. Increases in chromosome number occur typically by chromosome fission events, whereas decreases occur by translocation events that combine two large chromosomes and lead to a second smaller chromosome that is subsequently lost ^27^. Dysploidy is common, as indicated by chromosome numbers that deviate from euploid counts in numerous plant groups ^1, 2, 28, 29^. However, the relative contribution of dysploidy to chromosomal evolution alongside polyploidy is largely unexplored. Also, the effect of dysploidy on lineage diversification in the angiosperms remains broadly untested. Recent studies have suggested that dysploidy may elevate lineage diversification in focal plant clades, e.g., *Passiflora* ^30, 31^, Pooideae ^32^, and Cyperaceae ^30, 31^. However, a meta-analysis study ^33^ involving 15 plant clades (mostly genera or infrageneric sections) did not find a correlation between dysploid changes and lineage diversification.

Here, we explored variation in the rates of polyploidy and dysploidy across major angiosperm clades (orders and families). We examined the relationships between polyploidy and dysploidy with lineage diversification. To these ends, we combined chromosome numbers from the Chromosome Count Database (CCDB) ^2^ and a time-calibrated mega-phylogeny ^34, 35^ to assemble clade-wise data sets for 46 orders and 147 families of angiosperms. For each clade, we estimated the rates of polyploidy and dysploidy using ChromEvol ^36, 37^ and estimated the rate of net diversification independently of the rates of chromosome evolution using two methods: Magallón and Sanderson ^38, 39^ and Nee et al. ^38, 39^. We took this approach because of the lack of enough resolved phylogenies for the angiosperm orders and families examined here, which would have allowed us to relate chromosome evolution to speciation and extinction rates explicitly across a phylogeny. By analyzing the relationships between the rates of chromosomal evolution and the rate of net diversification across the major angiosperm groups, we provide taxonomically broad quantitative support that both modes of chromosomal evolution — polyploidy and dysploidy — likely play important roles in the formation of species and lineage radiation in the angiosperms.

## RESULTS

### Mode and pattern of chromosomal evolution across the angiosperms

Here, we performed a large-scale phylogenetic analysis of chromosome numbers of 46 orders and 147 families of angiosperms (**Supplementary Table 1**; **Supplementary Table 2**). In all, we analyzed 29,770 tip taxa (55% of them had chromosome counts) across the orders as well as 28,000 tip taxa (56% of them had chromosome counts) across the families. We found ChromEvol model support for polyploidization in all 46 angiosperm orders and 139 out of 147 families (**Supplementary Figure 1**; **Supplementary Table 3**; **Supplementary Table 4**). The eight families with no support for polyploidization were Alstroemeriaceae, Fagaceae, Grossulariaceae, Nepenthaceae, Nothofagaceae, Schisandraceae, Smilacaceae, and Tamaricaceae. The most common best-fitting model among the angiosperm orders was the four-parameter model “DEMI EST”, which allows for both polyploidization and demi-polyploidization, followed by the three-parameter model “DEMI”, which sets the rates of polyploidy and demi-polyploidization to be equal. Among the angiosperm families, the most common best-fitting model was “DEMI” rather than “DEMI EST”. Overall, in most clades, the best-fitting model allowed for both polyploidization and demi-polyploidization to occur.

Using ChromEvol, we reconstructed the pattern of chromosomal evolution across the angiosperm phylogeny under the best-fitting model. We found that the median gametic chromosome number at internal nodes was *n* = 11 across the major groups (**Supplementary Figure 2**), a number that was similar within each major group: in the asterids, *n* = 11; in the rosids, *n* = 11; in other (non-asterid/non-rosid) eudicots, *n* = 9; in the commelinid monocots, *n* = 13; in other (non-commelinid) monocots, *n* = 14; in the magnoliids, *n* = 12; in Austrobaileyales and Nymphaeales, *n* = 14. Moreover, the median ancestral chromosome number inferred at the root of the phylogeny was *n* = 7 for angiosperm orders and *n* = 9 for angiosperm families.

### Among-lineage variation in the rates of chromosomal evolution

The rates of polyploidy, as estimated under the four-parameter ChromEvol model, differed by 462 times and 447 times across the angiosperm orders and families, respectively (excluding eight families with a rate of nearly 0 EMY, i.e., < 1 x 10^-11^ EMY) (**Figure 1**; **Figure 2**; **Supplementary Figure 3**). The *order-wide* rate of polyploidy spanned from 0.00090 EMY in Arecales to 0.42 EMY in Poales (median, 0.024 EMY) (**Figure 1**). The *family-wide* rate of polyploidy ranged from nearly 0 EMY in eight families to 0.71 EMY in Poaceae (median, 0.025 EMY) (**Figure 2**; **Supplementary Figure 3**). We observed that the rosids underwent relatively high rates of polyploidy (12 out of 15 rosid orders had a rate above the median rate), and the non-commelinid monocots and the magnoliids experienced relatively low rates of polyploidy (one out of five non-commelinid monocot orders and none of the magnoliid orders had a rate above the median) (**Supplementary Table 5**; family-level results in **Supplementary Table 6**).

**Figure 1.**
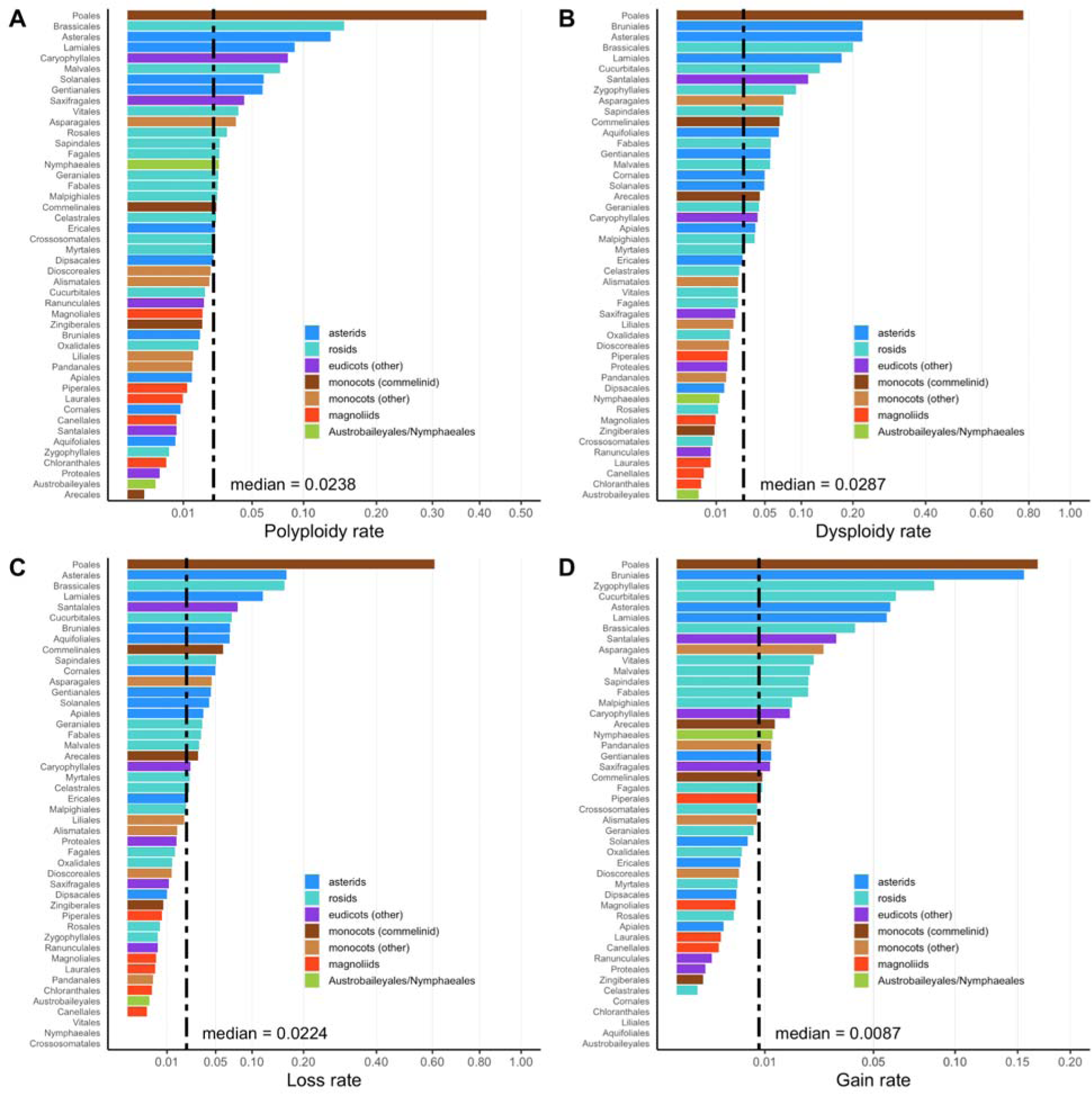
Angiosperm ***orders*** ranked by the rate of polyploidy (A), the rate of dysploidy (B), the rate of chromosome loss (C), and rate of chromosome gain (D) estimated under the four-parameter ChromEvol model. The x-axis scale is transformed by square root.

**Figure 2.**
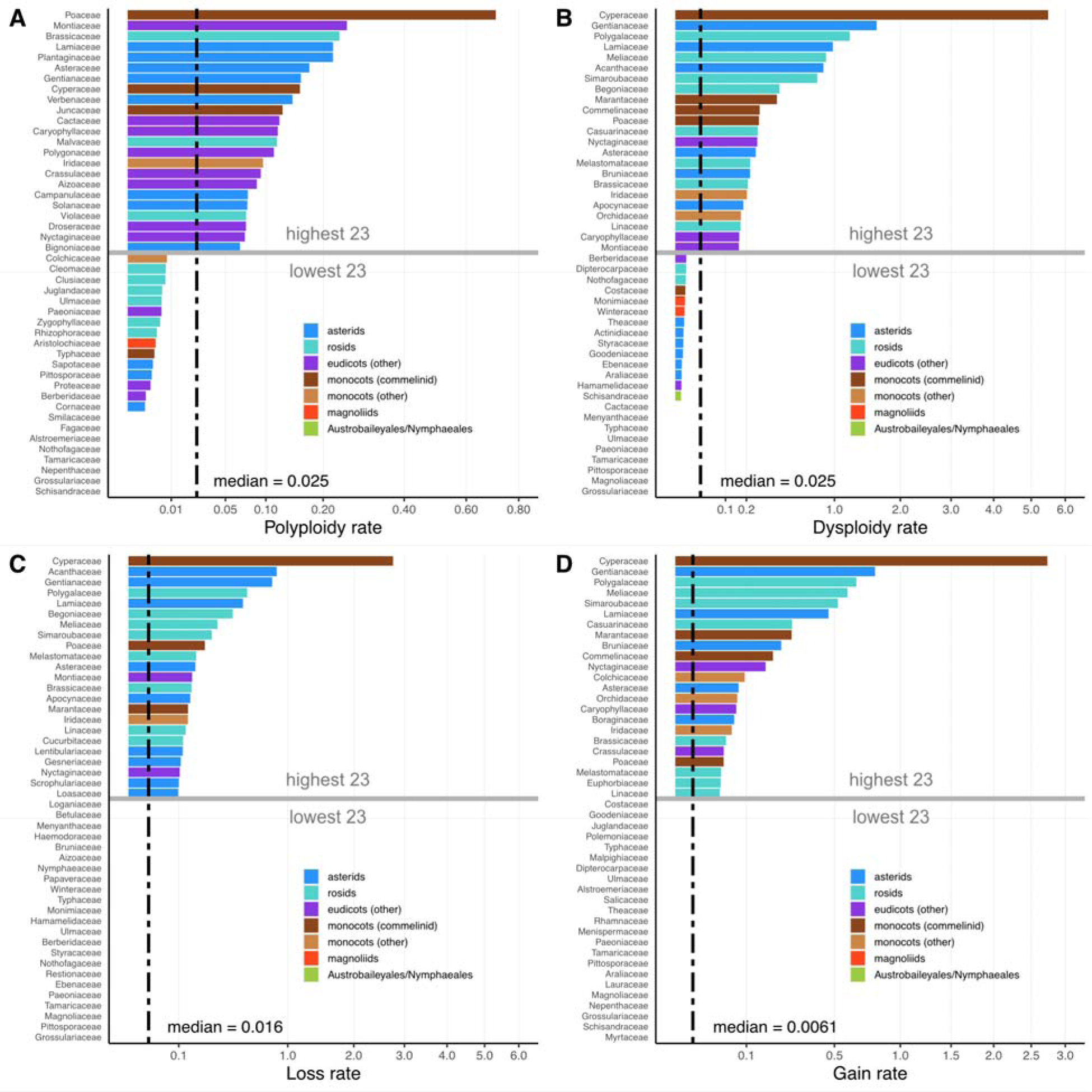
Angiosperm ***families*** ranked by the rate of polyploidy (A), the rate of dysploidy (B), the rate of chromosome loss (C), and the rate of chromosome gain (D) estimated under the four-parameter ChromEvol model. The x-axis scale is transformed by square root. Only the 23 clades with the highest rates and the 23 clades with the lowest rates for each type of rate are shown here due to space constraint (see **Supplementary Figure 3** for the full ranking).

Rates of dysploidy (the rate of chromosome loss plus the rate of chromosome gain), as estimated under the four-parameter ChromEvol model, were also remarkably variable. The rates of dysploidy differed by 251 and 4,205 times across the orders and families, respectively (excluding nine families with a rate of nearly 0 EMY) (**Figure 1**; **Figure 2**; **Supplementary Figure 3**). The *order-wide* rates ranged from 0.0031 EMY in Austrobaileyales to 0.78 EMY in Poales (median, 0.029 EMY) (**Figure 1**). The *family-wide* rate of dysploidy was as low as 0 EMY in nine families to as high as 5.50 EMY in Cyperaceae (median, 0.025 EMY) (**Figure 2**; **Supplementary Figure 3**). We noticed that the asterids underwent relatively high rates of dysploidy (eight out of 10 asterid orders had a rate above the median rate), and the non-commelinid monocots and the magnoliids experienced relatively low rates of dysploidy (one out of five non-commelinid monocot orders and none of the magnoliid orders had a rate above the median rate) (**Supplementary Table 5**; family-level results in **Supplementary Table 6**).

We next examined the rates of chromosome loss and gain separately (**Figure 1**; **Figure 2**; **Supplementary Figure 3**). The *order-wide* rates of loss ranged from nearly 0 EMY in three orders to 0.61 EMY in Poales (median, 0.022 EMY), whereas the rates of gain ranged from nearly 0 EMY in five orders to 0.17 EMY in Poales (median, 0.0087 EMY) (**Figure 1**). The *family-wide* rates of loss range from nearly 0 EMY in 29 families to 2.76 EMY in Cyperaceae, whereas the rates of gain range from nearly 0 EMY in 46 families to 2.73 EMY in Cyperaceae (**Figure 2**; **Supplementary Figure 3**). The asterids underwent relatively high rates of loss (nine of out 10 asterid orders had a rate above the median rate), and the non-commelinids and the magnoliids had relatively low rates of loss (one out of five non-commelinid orders and none of the magnoliid orders had a rate above the median rate) (**Supplementary Table 5**; family-level results in **Supplementary Table 6**); these trends were not observed with the rates of gain, except that the magnoliids experienced relatively low rates of gain (one out of five magnoliid orders had a rate above the median rate). These results indicate that generally, the rates of loss are higher than the rates of gain across the angiosperms.

Overall, the total rates of chromosomal evolution (the rate of dysploidy plus the rate of polyploidy) differed by 212 and 4,324 fold across the angiosperm orders and families, respectively (excluding two families with a total rate of nearly 0 EMY) (**Figure 3**; **Figure 4**; **Supplementary Figure 4**). Chromosomal evolution proceeded most slowly in Austrobaileyales (0.0056 EMY) and most rapidly in Poales (1.19 EMY; see below) (median, 0.053 EMY) (**Figure 3**). Generally, the rates of chromosomal evolution were lower in the monocots, the magnoliids, Austrobaileyales, and Nymphaeales (median, 0.016 EMY) and higher in the eudicots (median, 0.026 EMY) (p = 0.032, Wilcoxon’s test).

**Figure 3.**
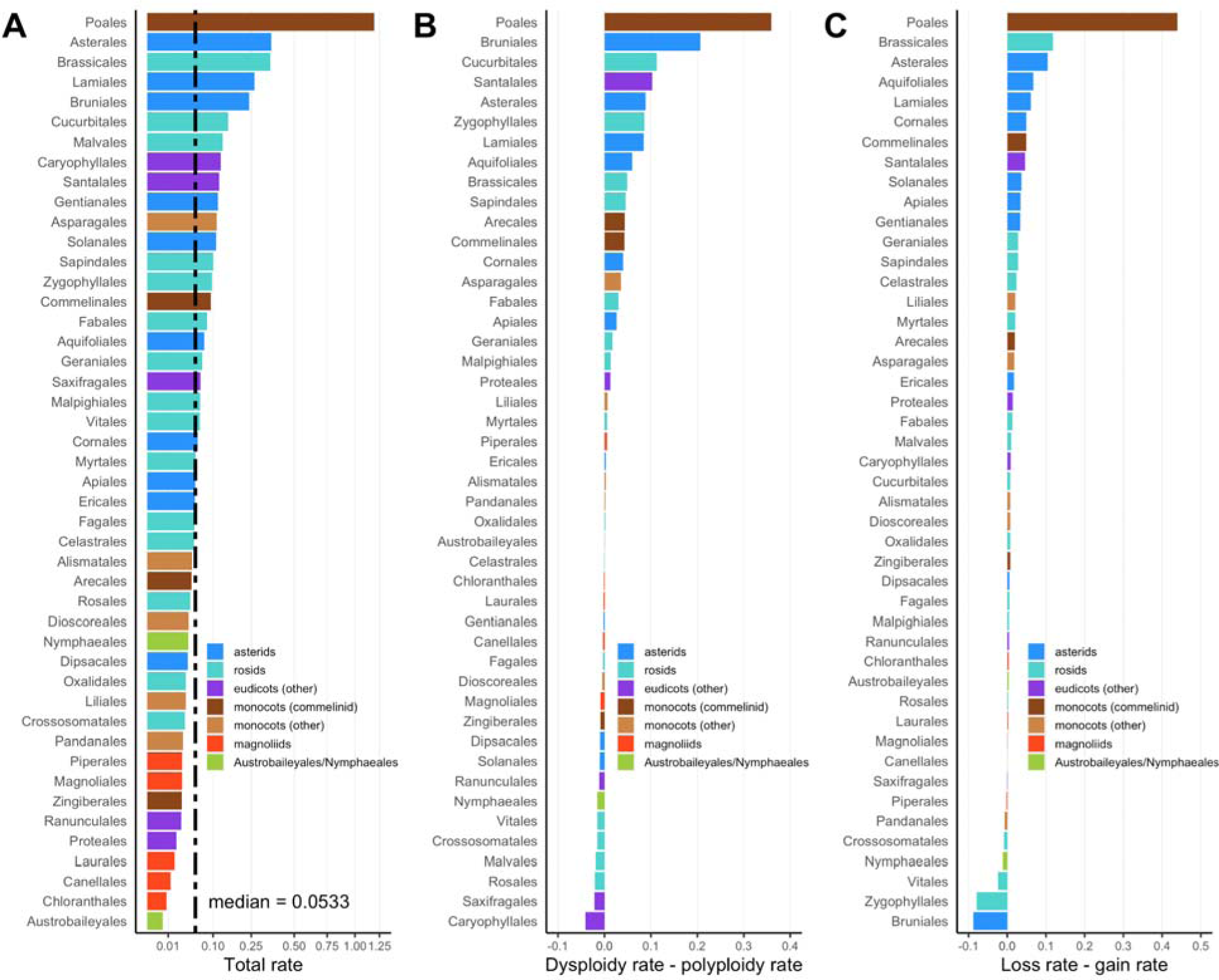
Angiosperm ***orders*** ranked by the total rate of chromosomal evolution (A), the difference between the rate of dysploidy and the rate of polyploidy (B), and the difference between the rate of chromosome loss and the rate of chromosome gain (C). The rates were estimated under the four-parameter ChromEvol model. The x-axis scale is transformed by square root.

**Figure 4.**
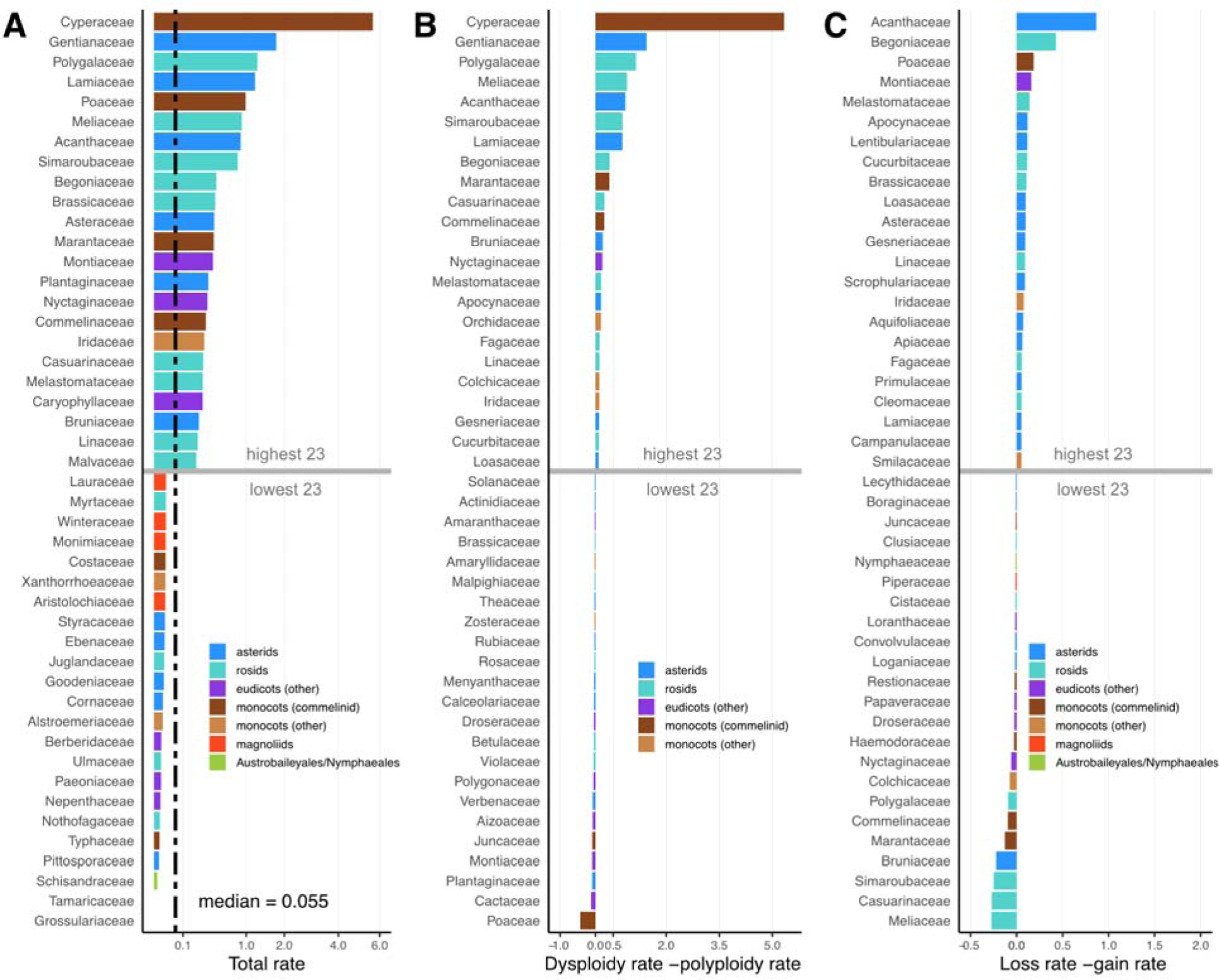
Angiosperm ***families*** ranked by the total rate of chromosomal evolution (A), the difference between the rate of dysploidy and the rate of polyploidy (B), and the difference between the rate of chromosome loss and the rate of chromosome gain (C). The rates were estimated under the four-parameter ChromEvol model. The x-axis scale is transformed by square root. Only the 23 clades with the highest rates and the 23 clades with the lowest rates for each type of rate are shown here due to space constraint (see **Supplementary Figure 4** for the full ranking).

We observed broad variation in the relative contribution of polyploidy and dysploidy to chromosomal evolution across the angiosperm clades. The rate of dysploidy was greater than the rate of polyploidy in 27 out of 46 (59%) angiosperm orders and in 69 out of 147 (47%) families (**Figure 3**; **Figure 4**; **Supplementary Figure 4**). This result suggests that dysploidy may be as pervasive as polyploidy throughout the evolutionary past of the angiosperms. Additionally, the rate of loss was greater than the rate of gain in 38 out of 46 (83%) angiosperm orders and in 98 out of 147 (67%) families (**Figure 3**; **Figure 4**; **Supplementary Figure 4**). This result suggests that chromosome loss is the predominant mode of dysploidy.

### Highest rates of chromosomal evolution in Poales

Poales displayed the most striking rates of dysploidy and polyploidy among the angiosperm orders (**Figure 1**) and was often an outlier in subsequent analyses. Two speciose families within Poales, the Cyperaceae and Poaceae, are known to experience frequent chromosomal changes ^23, 30^. Indeed, Cyperaceae experienced the highest total rate of chromosomal evolution (5.64 EMY) (median, 0.055 EMY), and Poaceae had the fifth highest total rate among the angiosperm families examined (**Figure 4**). Cyperaceae is well known for rampant dysploidy, especially in its largest genus, *Carex* ^40–42^. However, this family has a relatively low frequency of genome doubling except in non-*Carex* lineages ^43^. The exceptional rate of dysploidy in Cyperaceae is attributed to the holocentric chromosomes in many of its species ^43^. Indeed, our results found that dysploidy events were frequent in the Cyperaceae (5.5 EMY; 218 times the median rate of families) and polyploidy was estimated to occur at a lower rate than dysploidy (0.16 EMY; six times the median rate of families) (**Figure 2**; **Figure 4**). Conversely, we found that Poaceae experienced the highest rate of polyploidy among the angiosperm families (0.71 EMY; 28 times the median rate of families) but a lower rate of dysploidy (0.28 EMY; 11 times the median rate of families) (**Figure 2**); Poaceae still experienced a high total rate of chromosomal evolution (**Figure 4**). This result is consistent with previous observations that polyploidy is common in the grasses^23, 28, 44.^

### Net diversification rate is more correlated with polyploidy rate than dysploidy rate

It has long been posited that chromosomal rearrangements lead to reproductive isolation, thereby contributing to species formation and lineage diversification in the angiosperms ^45, 46^. To test for evidence of a relationship, we investigated whether the rates of chromosomal evolution (polyploidy and dysploidy) were correlated with the rate of net diversification. We found that the rates of net diversification estimated using MS method (assuming a relative extinction rate of 0.00, 0.50, or 0.90) and using the Nee method were highly positively correlated with each other (**Supplementary Figure 5**), and so we took the MS rate under an extinction rate of zero for correlation analysis with the rates of chromosomal evolution estimated under the four-parameter ChromEvol model. Simple correlation analyses indicated that the rates of polyploidy, dysploidy, and chromosomal evolution (polyploidy plus dysploidy) were positively correlated with the MS net diversification rates across angiosperm orders (**Supplementary Figure 6**; see **Supplementary Figure 7** for the results without Poales; **Supplementary Table 7**) and families (**Supplementary Figure 8**; see **Supplementary Figure 9** for the results without Cyperaceae and Poaceae; **Supplementary Table 8**). In the order-level analysis, the rate of chromosome loss – but not the rate of gain – was positively correlated with the MS net diversification rate (**Supplementary Figure 6**; **Supplementary Figure 7**); however, in the family-level analysis, both the rates of loss and gain were positively correlated with the MS net diversification rate (**Supplementary Figure 8**; **Supplementary Figure 9**). Patterns were obscured when Nee’s method was used to estimate diversification rates, likely because of uncertainty in extinction rates when using incompletely sampled phylogenies (**Supplementary Figures 6-9**).

Although the correlation analyses suggested a relationship between chromosomal evolution and diversification, the analyses were potentially confounded by shared error in a common variable. The rates of chromosomal evolution and net diversification both depend on the root age (in millions of years), which is estimated with error, like all phylogenetic age estimates. If the root age was under- or overestimated, that would elevate or diminish, respectively, both the rates of chromosome evolution and the rates of diversification, thereby artifactually leading to a positive correlation between the two. As an alternative approach, we examined the relative association between polyploidy versus dysploidy (or chromosome loss and gain) and net diversification, which did not suffer from the problem because the rates being compared (polyploidy versus dysploidy) were affected by the root age in the same way. We performed a regression analysis in conjunction with a relative importance analysis (RIA) to estimate the relative proportion of variance in the rates of net diversification explained by polyploidy and dysploidy. Our analysis indicated that the rate of polyploidy was more correlated with the MS net diversification rate than the rate of dysploidy in the angiosperm orders and the families (**Figure 5**). This trend was more apparent when analyzing the angiosperm families (polyploidy, 75% versus dysploidy, 25%) than the orders (polyploidy, 65% versus dysploidy, 35%) (**Figure 5**). Furthermore, when analyzing the rates of chromosome loss and gain separately (instead of a combined rate of dysploidy), an RIA revealed that the rate of polyploidy was more correlated with the MS net diversification rate than the rate of loss and the rate of gain in angiosperm orders (polyploidy, 56% versus loss, 35% versus gain, 10%) and families (polyploidy, 57% versus loss, 30% versus gain, 13%) (**Figure 5**). These qualitative results generally hold when the outlier clades (Poales, Cyperaceae, and Poaceae) were excluded (**Supplementary Figure 10**). Moreover, when the Nee net diversification rates were analyzed instead, the results of the RIA were generally reflected but were weaker, except in the order-level RIA considering the loss rate and gain rate separately (**Supplementary Figure 1****1**; **Supplementary Figure 1****2**).

**Figure 5.**
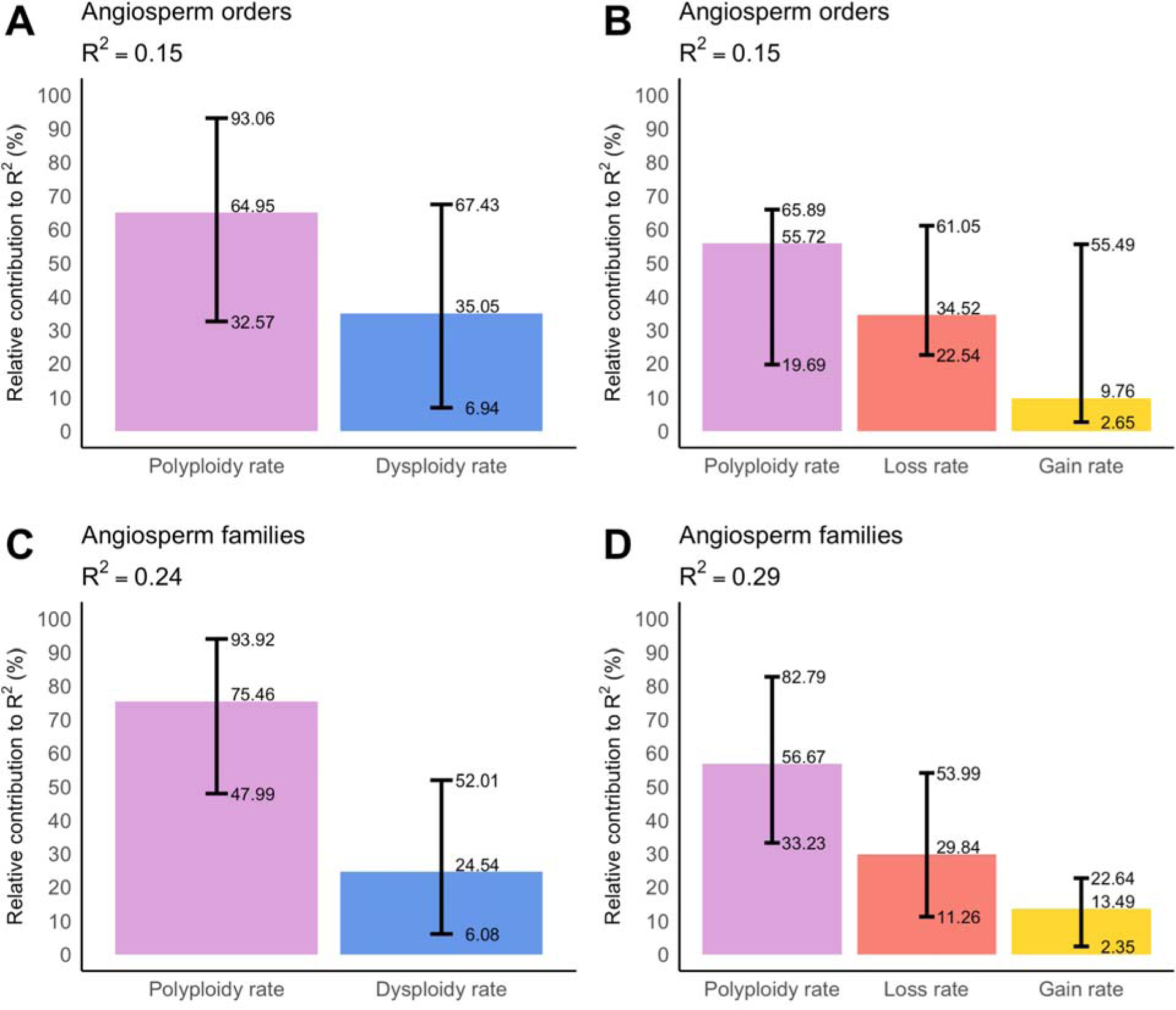
Relative association of the rates of chromosomal evolution to the rate of net diversification in angiosperm orders (A, B) and families (C, D). The net diversification rate was estimated as per Magallon and Sanderson (2011), setting the extinction rate to zero. Robust regression followed by a relative importance analysis was conducted using the method of Lindeman et al. (1980) with bootstrapping (1,000 replicates). The rates of chromosomal evolution were estimated under the four-parameter ChromEvol model. We considered two regression models; first we considered the rate of polyploidy and the rate of dysploidy (A, C), and second we considered the rate of polyploidy and the rates of chromosome loss and gain separately (B, D). The proportion of variance explained by each model is shown in the panel title. We estimated the relative contribution to R^2^ by the rates of polyploidy and dysploidy (A, C) and by the rates of polyploidy, loss, and gain (B, D). The bars represent 95% confidence intervals around the relative contributions.

## DISCUSSION

Our analyses of nearly 30,000 species found that the rates and modes of chromosomal evolution vary dramatically across the angiosperms. Consistent with previous research, we found that polyploidy ^47, 48^ and dysploidy are widespread and common ^33, 49^. Other recent studies of plant chromosomal evolution at this scale have capped chromosome numbers to keep computational time reasonable ^50^. In our analyses, we chose an alternative approach and analyzed individual families and orders without excluding species with higher chromosome numbers. This allowed us to compare and rank variation among angiosperm clades. We found that the rates of chromosomal evolution were generally lower in the monocots, the magnoliids, Austrobaileyales, and Nymphaeales (median, 0.016 EMY) and higher in the eudicots (median, 0.026 EMY) (p = 0.032, Wilcoxon’s test). Although many features may explain this trend, growth form (i.e., herbaceous versus woody lineages) likely underlies some of the variation in the rate of chromosomal evolution ^51^. The magnoliid orders Laurales, Canellales, and Chloranthales exhibit some of the lowest rates of polyploidy and dysploidy, along with Austrobaileyales. These four orders are predominantly woody (i.e., trees and shrubs). Intriguingly, the remaining magnoliid order Piperales contains an assortment of herbaceous and woody plants and has a higher rate of dysploidy. On the other end of the spectrum, the monocots Poales and Commelinales and the dicots Asterales, Brassicales, and Bruniales are herbaceous plants with the most dynamic karyotypes among the orders examined, undergoing the highest rates of polyploidy and dysploidy. These results are consistent with the finding of a recent study ^52^ showing the association between polyploidy and growth forms (higher frequency of polyploidy in herbaceous versus woody plants), a hypothesis proposed early on by Stebbins ^53^. Moreover, it is quite possible that polyploidy is associated with other traits yet to be examined jointly with polyploidy ^16, 26, 54^. For example, Stebbins ^53^ observed that polyploidization occurs more frequently in annual and biennial lineages than in perennial lineages. Combined analyses of traits and chromosomal evolution are needed to better understand what underlies the impressive variation in the rates of polyploidy and dysploidy across the angiosperms.

Looking more closely at polyploidy, we found that polyploidization rates varied over 400-fold across families and orders of angiosperms. Although previous research has indicated that the frequency and incidence of polyploidy varies among angiosperm clades ^17, 47, 48^, prior studies have not comprehensively quantified this dramatic difference in the rates of polyploidy. Polyploidy was most frequent in the Poaceae with an estimated polyploid speciation event every 0.71 million years, consistent with expectations of the frequency of polyploidy in the family ^1, 55^. The rate of polyploidization in the Poaceae was nearly three times the rate of the next two families, the Montiaceae and the Brassicaceae. Overall, the estimated rates of polyploidization were similar to estimates of the rate of plant speciation from other phylogenetic analyses ^56^. Perhaps most surprising given the association of polyploidy with flowering plant evolution is that we detected no polyploidy in eight families. To provide some external validation of our analyses with different data and confirm if polyploidy is rare or absent in these families, we compared our results to inferences of ancient WGDs in the 1KP data set ^15, 57^. We may expect that if recent polyploidy is generally rare in these families they may not have any signs of ancient polyploidy either. Six of the families were represented in the 1KP study (Fagaceae, Grossulariaceae, Nepenthaceae, Nothofagaceae, Schisandraceae, and Smilacaceae). There was no evidence for ancient polyploidization in three (Fagaceae, Grossulariaceae, and Nothofagaceae), but there is evidence for WGD in the ancestry of Nepenthaceae (NEPNα), Schisandraceae (ILFLα), and Smilacaceae (SMBOα) ^15, 57^. The sampled contemporary chromosome numbers in Nepenthaceae and Schisandraceae did not indicate recent polyploidization. However, because ChromEvol models only chromosome counts — a gross feature of genome organization — it may not be able to detect ancient WGDs in lineages that have fully diploidized. In Smilacaceae, there is one possible instance of recent polyploidization (*n* = 48; *n* = 13 to 16 in other taxa), but it was not enough for model support for polyploidization. Beyond these exceptional families, polyploidy was a major feature of nearly all families and occurred frequently throughout the history of flowering plants.

Rates of dysploidy varied more than 4,000-fold across flowering plant families, an order of magnitude more than polyploidy. Perhaps unsurprisingly, this variation was driven by rapid rates of dysploidy in the holocentric Cyperaceae ^30^. We estimated that nearly 5.5 dysploid changes occur million years in Cyperaceae, more than three times higher than the next fastest family, the Gentianaceae. Holocentric chromosomes are generally thought to drive high rates of chromosomal evolution ^58^, but a recent study in insects found that holocentric lineages did not have higher rates of chromosomal evolution than monocentric lineages ^59^. Similarly, besides Cyperaceae, other families with at least some holocentric taxa ^58^ did not have exceptional rates of dysploidy in our analyses. More detailed analyses are needed to assess if the holocentric lineages in these families do have higher rates of chromosomal evolution, and rigorously test if holocentricity is a major driver of dysploidy in plants.

Overall, we found that chromosome loss was the predominant form of dysploidy, as found in other recent research ^50^. Rates of loss were higher than the rates of gain across most clades of angiosperms (**Figure 3**). The predominance of descending dysploidy is important because chromosome numbers across flowering plants occur in a relatively tight range centered around *n* = 7–12 ^2, 60^ despite multiple rounds of polyploidy ^4, 15, 57^. Most flowering plants are estimated to have experienced 4 to 6 rounds of WGD in their ancestry ^15^. Descending dysploidy following polyploidy must have repeatedly brought these numbers back down to this range. In our estimates, the rates of dysploidy and polyploidy were similar in many families and orders.

However, there is evidence that chromosome numbers may be reduced rapidly following polyploidy ^61–66^. This could be problematic for estimating rates in ChromEvol because the models assume that the rate is equally distributed across branches. Implementing models that allow additional parameters or approaches to account for the likely heterogeneity of dysploidy, especially in concert with WGDs, may provide better rate estimates. Regardless, the root numbers estimated in our analyses were consistent with the analysis of Carta et al. ^50^ despite different approaches and scales of data, and this number, *n* = 7, has long been suggested as a base number for flowering plants ^1, 28^.

The changes driving these genome dynamics are also thought to drive the diversity of flowering plants. Our analyses uncovered evidence for a positive relationship between chromosomal evolution and diversification. Net diversification rates were positively correlated with the rates of polyploidy and dysploidy in the angiosperms. Combining both types of chromosome changes, Levin and Wilson ^51^ observed a similar positive correlation in the seed plants (rate of overall karyotypic change versus rate of speciation), despite that their pioneering analysis used different methods and data. In a meta-analysis, Escudero et al. ^33^ found a lack of correlation between dysploid changes and lineage diversification in 15 plant genera or infra-generic groups.

However, there is a large difference in the evolutionary time scales considered. Escudero et al. ^33^ tested the relationship at relatively short evolutionary time scales (since the divergence of genera or groups nested therein). Our study, on the other hand, examined the long-term rates and consequences of dysploid changes on lineage diversification (since the divergence of orders and families). Consistent with our results, Márquez-Corro et al. ^30^ reported that dysploidy is associated with increased net diversification in Cyperaceae. Similarly, considerable differences in the evolutionary time scales may help explain the seemingly contradictory results that polyploid lineages are correlated with lower net diversification rates within angiosperm genera ^17^ versus higher diversification across the deep evolutionary past of the angiosperms ^13, 14^. Many polyploid lineages may be doomed to extinction over the short term, for example, but those that do persist may be a biased subset that can persist and eventually benefit from the duplicated genes.

Using RIA, we teased apart the relative contributions of the rates of polyploidy and dysploidy to the rate of net diversification. We found that the rate of polyploidy explained more of the variation in net diversification than the rate of dysploidy in the angiosperms. It is hypothesized that dysploidy may have a lesser effect on lineage diversification than polyploidy ^33, 67^, because dysploidy generally neither alters the DNA content nor disrupts gene dosage balance, whereas polyploidy tends to exert noticeable effects on an organism’s phenotype. Indeed, polyploids have been associated with physiological changes (e.g., heterosis, ^68^; changes in secondary metabolite content ^69–71^; drought tolerance, ^72, 73^; reviewed in ^8^) and niche differentiation and ecological changes ^6, 8, 11, 74, 75^, although nascent polyploids often suffer from negative effects (such as reduced hybrid fertility ^19^) and inefficient selection when genes are masked by multiple copies ^20–22^. Despite the observation that many polyploid lineages may be evolutionary “dead-ends” in the short term ^17^, they enjoy occasional moments of success ^9, 14, 15, 25^. Furthermore, polyploidy has been found to interact with other traits that shape the outcome of polyploid success ^16, 26^ and may explain the complicated relationship between polyploidy and diversification.

There are two reasons that polyploidy may be correlated with higher net diversification rate, which cannot be distinguished with the correlation analyses used here. One explanation is that polyploid or dysploid species diversify at a higher rate than other species, but a second explanation is that diploids that readily undergo polyploidization have access to a faster route to speciation than diploids that rarely undergo polyploidization, leading to a correlation between groups with more species and groups with more polyploids. The same applies to dysploid routes to speciation. Previous work ^16, 17, 76^ has found stronger evidence for the second explanation than the former. Future analyses, such as the phylogenetic path analyses in Roman-Palacios et al. ^16^, are needed to tease apart these mechanisms.

Our results suggest that dysploid changes may nevertheless have effects on cladogenesis and lineage diversification in the angiosperms that have been largely overlooked. Others have posited that post-polyploidization diploidization (via chromosome number reduction alongside genome downsizing) could lead to lineage radiation ^62, 64, 65, 77^. In support of this hypothesis, our results indicate that the rate of chromosomal loss is better correlated with the rate of net diversification than the rate of gain. Also, an intriguing, but complicating, possibility may be that the effects of dysploid changes on diversification are delayed ^62^, in a vein similar to the “lag-time” hypothesis proposed for ancient polyploidization events in plants ^13, 78^. Detailed studies of particular groups (e.g., ^63, 64^) would be insightful to disentangle the impact of polyploidy and dysploidy on diversification in the angiosperms.

Dysploid changes arise from mechanistic errors during meiosis, which produce gametes with unequal sets of chromosomes. The connection between dysploid changes and diversification have been investigated in lineages with holocentric chromosomes, particularly Cyperaceae (e.g., Márquez-Corro et al. ^30^). Márquez-Corro et al. ^79^ reviewed existing data for holocentric lineages and their monocentric kins across the eukaryotes. The workers found that differences in chromosome pairing behavior during meiosis may be associated with differences in lineage diversification. Data for holocentricity across the angiosperm lineages are scarce, however. Only a few plant clades are known to have holocentric species ^79^. Nevertheless, it would be fruitful for future research to disentangle the intricate relationship between holocentricity, meiotic chromosome pairing behavior, and diversification in other angiosperm groups, such as Marantaceae and Gentianaceae, that also experience frequent dysploidy.

The contribution of polyploidy to speciation and lineage diversification in the angiosperms has been widely appreciated, but dysploidy has received far less attention. Our results indicate that variation in the rates of polyploidy and dysploidy are positively correlated with the rate of net diversification in the angiosperms. In turn, the among-lineage disparity in the rates of chromosomal evolution may be attributed to differences in centromere structure ^30^, variation in the mechanisms of meiotic chromosome pairing ^66^, growth form and life history ^51–53^, mating system ^23, 26, 54^, and other unrecognized traits ^16^. A profitable direction of future studies may be more sophisticated joint phylogenetic modelling of chromosome number change, trait evolution (e.g., growth form and life history), and diversification dynamics (see ^16, 80, 81^). This interplay of karyotypic, genomic, and phenotypic features is undoubtedly complex; how this interplay sways the fates of angiosperm lineages remains a rich area for future research.

## MATERIALS AND METHODS

### Data curation

We collated (1) the phylogenetic trees, (2) chromosome numbers, and (3) estimates of species richness of 46 angiosperm orders and 147 families by combining data from several sources.

First, we prepared the phylogenetic trees of major angiosperm clades. Subtrees corresponding to angiosperm orders and families were extracted from the time-calibrated mega-phylogeny of Qian and Jin ^35^ (obtained from their online Supplementary Data), using the node labels made by the authors as a guide. The mega-phylogeny of Qian and Jin ^35^ is a corrected and expanded version of the mega-phylogeny originally reconstructed by Zanne et al. ^34^. We checked whether there were any polytomies in the subtrees that might interfere with our analyses. We encountered one polytomy in the Brassicales tree; it was randomly resolved once, and the length of the new branch was set arbitrarily to 10^-8^. The mega-phylogeny and its subtrees were processed using the R package *ape* version 4.1 ^82^.

Next, we obtained the chromosome numbers of the angiosperm clades for which phylogenies were available. Chromosome numbers were downloaded from the CCDB version 1.45 ^2^ (accessed on Oct. 12, 2017) For many species, the CCDB provides multiple entries of gametic or somatic chromosome counts. To summarize the chromosome numbers for a single species, the mode gametic chromosome number (*n*) was taken as the representative count. The mode count most likely captures the putative base number and diploid cytotype, which is the most common cytotype ^47^; it also minimizes the influence of unusual individual karyotypes. Somatic counts (2*n*) were halved and then rounded down before they were pooled together with the gametic counts to determine the mode count. Also, in instances where multiple mode counts occur for a species, the lowest mode count was retained, because it more likely reflects the diploid cytotype of the species (e.g., the mode count is *n* = 4 in this hypothetical series *n* = 4, 4, 8, 8, 16). The representative species mode counts were matched to the tips of the phylogenetic trees according to their binomial names. Before that, synonyms and spelling errors in the taxon names of the phylogenies were resolved using the same approach taken by Rice et al. ^2^. Briefly, the taxon names were matched to the accepted species names in The Plant List version 1.1 ^83^ using Taxonome version 1.5 ^84^. This step facilitated matching of the chromosome numbers with the tips of the phylogenetic trees, as the species names in the Qian and Jin ^35^ mega-phylogeny were also standardized to The Plant List. Only clades having more than 10 tip taxa with matched chromosome counts were retained for analysis.

Lastly, we recorded the number of extant species of each angiosperm clade from the Angiosperm Phylogeny Website (APW) version 14 ^85^ (accessed on Oct. 4, 2017). For one order and five families, a range of species numbers was available; in these instances, the midpoint was taken as the species number of the clade. For example, the APW indicated the species number of Amaranthaceae ranges from 2,050 to 2,500; we took 2,275 to be the species richness of this family. Also, the main estimate of species richness was taken whenever there were alternative estimates.

The above data curation procedure resulted in 46 order-level and 147 family-level angiosperm data sets. There were 29,770 tip taxa in total in the angiosperm order phylogenies, and 16,331 (55%) of these taxa had chromosome counts. On average, the percentage of taxon sampling (number of tip taxa divided by the species count from APW) in the order phylogenies was 18% (range, 3% to 75%), and the percentage of the tip taxa with chromosome counts in those phylogenies was 52% (range, 22% to 87%) (**Supplementary Table 1**). Also, there were 28,000 tip taxa in total in the angiosperm family phylogenies, and 15,809 (56%) of these taxa had chromosome counts. On average, the percentage of taxon sampling in the family phylogenies was 21% (range, 2% to 100%), and the percentage of the tip taxa with chromosome counts in the phylogenies was 57% (range, 10% to 97%) (**Supplementary Table 2**).

### Estimation of clade-wide rates of chromosomal evolution

Using ChromEvol ^36, 37^, we estimated the clade-wide rates of polyploidy and dysploidy for each angiosperm order and family. The ChromEvol models describe chromosome number change as a continuous-time Markov process along a phylogeny with observed chromosome numbers assigned to the tip taxa. ChromEvol jointly estimates the rates of chromosome gain (i.e., single increment), loss (i.e., single decrement), polyploidy (i.e., doubling), and demi-polyploidy (i.e., multiplication by 1.5 or half-doubling, e.g., transition between tetraploids and hexaploids) in a maximum likelihood framework.

Here, we applied four basic ChromEvol models that assume that the rate of chromosome gain (λ), the rate of chromosome loss (δ), the rate of polyploidy (ρ), and the rate of demi-polyploidy (μ) are constant throughout a given phylogeny. In the “CONST RATE” model, μ is fixed to zero while λ, δ, and ρ are allowed to vary; in “DEMI”, ρ is set to equal to μ while λ and δ are allowed to vary; in “DEMI EST”, all four rate parameters are allowed to vary; and in “NO DUPL”, ρ and μ are both set to zero while λ and δ are allowed to vary (therefore, no polyploidization occurs).

We infer that there is no evidence for polyploidy if no model is favoured over the “NO DUPL” model.

ChromEvol was run on the default optimization scheme, while setting the minimum allowed chromosome number to one and the maximum allowed chromosome number to be twice the highest chromosome number observed in a given dataset (e.g., in Malvaceae, the highest species gametic mode count is *n* = 135, and therefore the maximum allowed chromosome number was set to *n* = 270). The upper limit was applied to ensure that the transition rate matrix is reasonably large to capture a plausible range of chromosome numbers (i.e., allowing doubling for the species with the highest chromosome number), while maintaining computational tractability. In one instance, the gametic count of *Voanioala gerardii* (Arecaceae, Arecales) was exceptionally large (*n* = 298); the ChromEvol runs for Arecaceae and Arecales hardly proceeded even after a month, and so we reran ChromEvol by treating the chromosome count of *V. gerardii* as missing. For each clade, we determined the best-fitting ChromEvol model to be the one with the lowest Akaike Information Criterion (AIC) ^86^.

We considered the clade-wide rates of polyploidy (defined as the sum of the rate of polyploidy and the rate of demi-polyploidy) and dysploidy (defined as the sum of the rate of chromosome loss and the rate of gain) under each fitted ChromEvol model. We found that the rates of polyploidy and the rates of dysploidy were highly similar when estimated under the four-parameter model (i.e., “DEMI EST”) and the best-fitting model, which might not necessarily be the four-parameter model. We used the results obtained under the best-fitting models to assess ChromEvol model support across the data sets (**Supplementary Figure 1**) and to reconstruct ancestral chromosome numbers in the phylogenies (**Supplementary Figure 2**). When we analyzed the rate estimates, we took the results obtained under the four-parameter model to avoid biases in the rate estimates that could be introduced by forcing the rate of polyploidy and/or the rate of demi-polyploidization to be zero. The results of the analyses using the rates estimated under the four-parameter model are presented in **Figures 1-5** and **Supplementary Figures 3, 4, 6-12**.

### Estimation of clade-wide rates of net diversification

To estimate the clade-wide net diversification rate (speciation rate minus extinction rate) of each angiosperm order and family, we employed two approaches: Magallón and Sanderson ^38, 39^ and Nee et al. ^38, 39^.

Magallón and Sanderson ^38, 39^, or MS, proposed a simple method to estimate the absolute diversification rate of a clade. The MS method assumes a pure birth model, leading to exponential growth, so that the rate of lineage diversification of a clade is log(*n*)/*t*, where *n* is its number of extant species in the clade and *t* is the age of the clade. For each clade, we set *n* to be the species richness obtained from the APW and *t* to be the crown age taken from the mega-phylogeny of Qian and Jin ^35^. Magallon and Sanderson ^38^ also suggested a means to correct for the number of unobserved taxa due to extinction. This involves assuming a relative extinction fraction, or the extinction rate divided by the speciation rate. The standard practice is to set the relative extinction fraction to different values (e.g., 0.00, 0.50, and 0.90 in Scholl and Wiens ^87^) and then to estimate the net diversification rate under each value. We observed that the different values (0, 0.50, and 0.90) gave estimates that were highly correlated (r > 0.98; Pearson’s correlation). Hence, we took the net diversification rates computed assuming an extinction rate of zero in later correlative analyses. The rates were calculated using the implementation of the MS method in the R package *geiger* version 2.0.6 ^88^.

Next, we estimated net diversification rates using the constant-rate birth-death model of Nee et al. ^39^. The Nee model describes the rates of speciation and extinction as functions of the branching time distribution of a phylogeny. The rates of speciation and extinction were inferred using a maximum likelihood approach, which was implemented using the R package *diversitree* version 0.9-10 ^89^. We accounted for random, incomplete taxon sampling by providing an estimate of sampling fraction (i.e., the number of tip taxa divided by the species count collected from the APW). Due to numerical imprecision from rounding, the subtrees appeared to be non-ultrametric; that is, the variation among root-to-tip distances within each tree exceeded the default machine tolerance used by *diversitree*. Therefore, we made minuscule corrections to the branch lengths of the subtrees so that the subtrees were ultrametric. We made these corrections using the non-negative least-squares method, as implemented in the R package *phangorn* version 2.4.0 ^90^ via *phytools* version 0.6-60 ^91^.

We conducted a maximum likelihood (ML) procedure as follows. To reduce the possibility of the ML search getting stuck at local optima, we ran the search ten times using ten starting points for the rates of speciation and extinction, which were randomly drawn from exponential distributions (whose means were equal to the rates of speciation and extinction heuristically estimated under the Nee model). The ML search was conducted using the subplex optimization algorithm ^92^ implemented in the R package *subplex* version 1.5-4 ^93^. For each clade, we took the net diversification rate that had the highest likelihood score to be the ML estimate.

The above analyses yielded estimates of the net diversification rate (events per million years, or EMY) obtained by two methods: (1) MS assuming an extinction rate of zero and (2) Nee under ML. These rates of net diversification are positively correlated with each other (**Supplementary Figure 3**). In the main results, we focus on the rates estimated under the simpler method, that is, MS assuming an extinction rate of zero. Moreover, the MS estimator is useful for poorly sampled clades (in our dataset, 26% and 27% of the angiosperm orders and families had a sampling fraction of at most 10%, respectively), because it does not require a well sampled species-level phylogeny, which is needed by approaches that require branch length information such as that of Nee et al. ^39^.

### Robust regression combined with relative importance analysis

A standard multiple linear regression analysis would not be meaningful here for two reasons. First, there are clearly influential outliers (Poales, Cyperaceae, and Poaceae), which experience exceptionally high rates of chromosomal evolution (**Figure 1**; **Figure 2**). Because those outliers are genuine biological anomalies, we wished to include them in our analyses, downweighting their importance as described below (see **Supplementary Figures 5, 7, 8, 10** for analyses where these clades were excluded). Second, the rates of polyploidy and dysploidy exhibit collinearity. Indeed, Pearson’s correlation tests indicated that the rate of polyploidy and the rate of dysploidy are positively correlated (orders, r = 0.91, p = 2.2 x 10^-16^; orders except Poales, r = 0.60, p = 1.3 x 10^-5^; families, r = 0.23, p = 0.005; families except Cyperaceae and Poaceae, r = 0.33, p = 6.1 x 10^-5^). To address these two issues, we performed robust regression coupled with a relative importance analysis.

Robust regression handles outliers by down-weighting them as per their influence. Here, we applied the method of M-estimation with Huber weighting ^94^, as implemented in the R package *MASS* version 7.3-50 ^95^. Next, we conducted a relative importance analysis, taking as input the model fit by M-estimation, to estimate the proportion of variance in the rate of net diversification explained by differences in rates of polyploidy and dysploidy. We obtained the relative contributions of polyploidy and dysploidy using the recommended method of Lindeman et al. ^96^. Briefly, the Lindeman et al. ^96^ method computes the relative proportions of the explained variance (i.e., the decomposed R^2^ of each predictor variable divided by the total R^2^) averaged over all possible sequential orderings of the independent variables, rather than basing the relative proportions on a single arbitrarily chosen ordering. We performed bootstrapping (1,000 replicates) to obtain 95% confidence intervals around the relative contributions of the rates of polyploidy and dysploidy to the rate of net diversification. We repeated this analysis for the MS rate (assuming an extinction rate of zero) and the Nee rate. The relative importance analyses were done using the R package *relaimpo* version 2.2-3 ^97, 98^.

## ACKNOWLEDGMENTS

Foremost, we thank the cytology community for its data collection effort over the past century. We thank Sean W. Graham, Wayne P. Maddison, Itay Mayrose, Emma E. Goldberg, Rosana Zenil-Ferguson, and members of the Otto lab and Barker lab for their insightful comments; Michal Drori for helping to parse the CCDB data; Anthony Baniaga for help with obtaining species richness; and Zheng Li for helping with navigating the 1KP data. This study was facilitated by the computing resources of Fusion Genomics Corp., Compute Canada, the University of British Columbia, and the University of Arizona. SHZ was supported by the UBC Four Year Doctoral Fellowship, the UBC Bioinformatics Training Program, an CIHR Doctoral Research Award, and an NSERC grant (RGPIN-2016-03711) awarded to SPO (UBC).

## AUTHOR CONTRIBUTIONS

SHZ designed and conducted the study, wrote the original draft of the manuscript, and reviewed and edited the manuscript. SPO provided supervision and reviewed and edited the manuscript. MSB conceived of and designed study, provided supervision, and reviewed and edited the manuscript.

## COMPETING INTERESTS

The authors have no competing interests to declare.

## SUPPLEMENTARY MATERIALS

**Supplementary Table 1.** Information about the angiosperm ***orders*** examined in this study. The order-level phylogenies were extracted from the Qian and Jin (2016) mega-phylogeny. The crown ages of the orders (in millions of years, or MY) were taken from these phylogenies and the species counts of the orders were taken from the Angiosperm Phylogeny Website (APW). For each clade, the number of tip taxa in the phylogeny, the number of tip taxa having chromosome counts, the taxon sampling fraction (number of tip taxa in the phylogeny divided by the species count), and the chromosome count sampling fraction (number of tip taxa with chromosome count divided by the number of tip taxa in the phylogeny) are presented.

**Supplementary Table 2.** Information about the angiosperm ***families*** examined in this study. See the explanation in **Supplementary Table 1.**

**Supplementary Table 3.** Results of the ChromEvol analysis for the angiosperm ***order***-level data sets. The best-fitting model (out of “NO DUPL”, “CONST RATE”, “DEMI”, and “DEMI EST”; see **Materials and Methods** for a description of the models) was inferred based on the lowest AIC score. The rates of chromosomal evolution (in events per a million years, or EMY) estimated under the best-fitting model and the full four-parameter model (“DEMI EST”) are provided. The rate of dysploidy was defined as the sum of the rate of chromosome loss (-1) and the rate of gain (+1). The rate of polyploidy was defined as the sum of the rate of duplication (2x) and the rate of demi-duplication (1.5x). The ancestral gametic chromosome number was inferred at the root of each clade under the best-fitting and four-parameter model.

**Supplementary Table 4.** Results of the ChromEvol analysis for the angiosperm ***family***-level data sets. See the explanation for **Supplementary Table 3.**

**Supplementary Table 5.** Number of angiosperm ***orders*** that have rates of chromosome evolution above the median rate in each major angiosperm group. Total rate was defined as the sum of the rate of polyploidy and the rate of dysploidy, and the dysploidy rate was defined as the sum of the rate of loss and the rate of gain. The bottom row indicates the number of clades in each major angiosperm group. For example, there were 7 asterid orders that had a total rate above the median rate of the 10 angiosperm orders examined here.

**Supplementary Table 6.** Number of angiosperm ***families*** which have rates of chromosome evolution above the median rate in each major angiosperm group. See the explanation for **Supplementary Table 5.**

**Supplementary Table 7.** Results of the diversification analysis for the angiosperm ***order***-level data sets. Absolute diversification rates were estimated using the method of Magallon and Sanderson (2001) assuming a relative extinction fraction of 0.00, 0.50, or 0.90. Additionally, net diversification rates were estimated using the method of Nee et al. (1994) under maximum likelihood, which infers the speciation rate and extinction rate of a given phylogeny.

**Supplementary Table 8.** Results of diversification analysis for the angiosperm ***family***-level data sets. See the explanation for **Supplementary Table 7.**

**Supplementary Figure 1.**
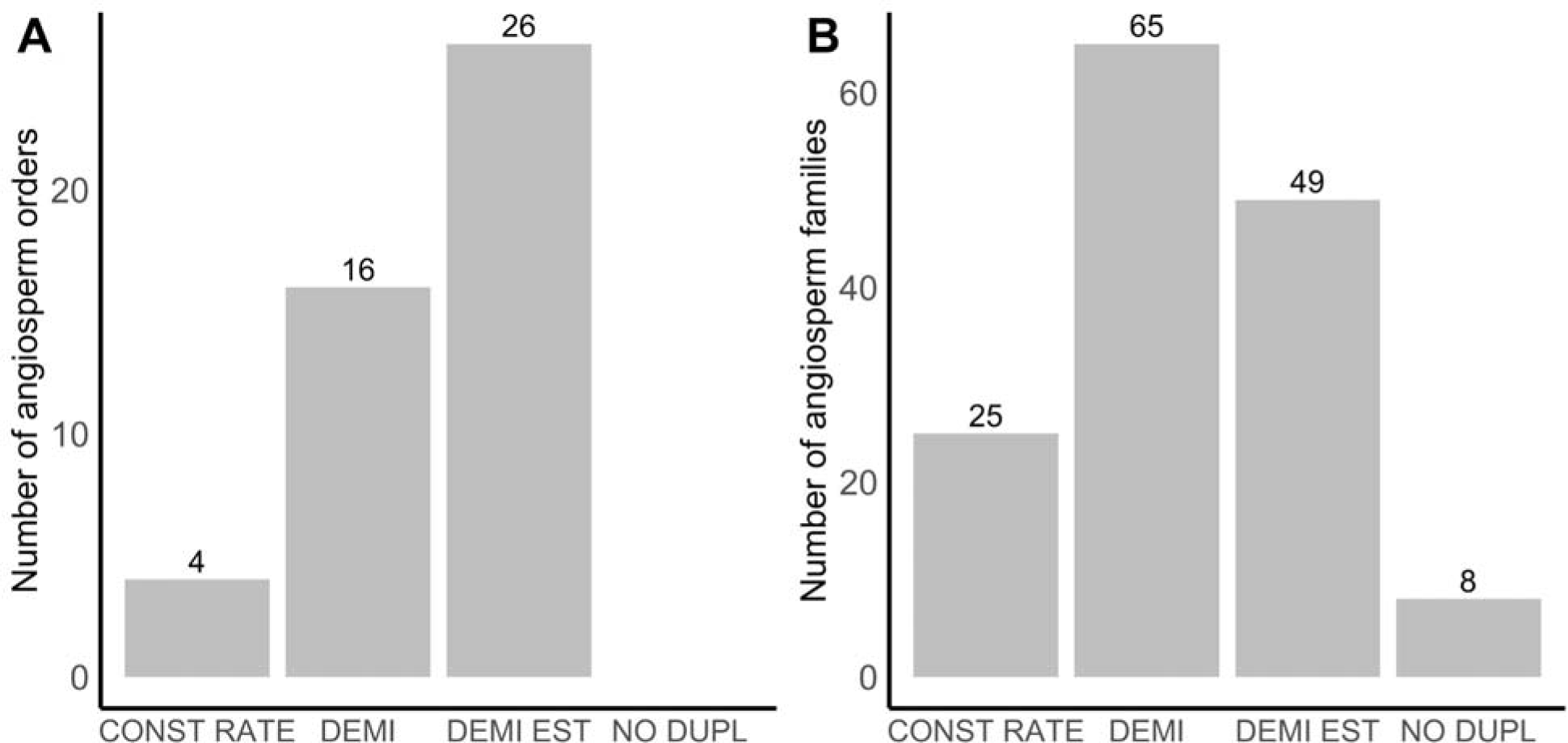
Best-fitting ChromEvol models for the angiosperm ***orders*** (A) and ***families*** (B) examined in this study. The models have four rate parameters: (1) chromosome λ; (2) chromosome loss or descending dysploidy, δ; (3) doubling or polyploidization, ρ and (4) half-doubling or demi-polyploidization, μ In the “CONST RATE” model, μ is fixed to zero; in “DEMI”, ρ is set to equal to μ in “DEMI EST”, all four rates are allowed to vary; and in “NO DUPL”, ρ and μ are both set to zero (therefore, no polyploidization is allowed). In all the angiosperm orders and most of the families (except eight cases), there was model support for polyploidization.

**Supplementary Figure 2.**
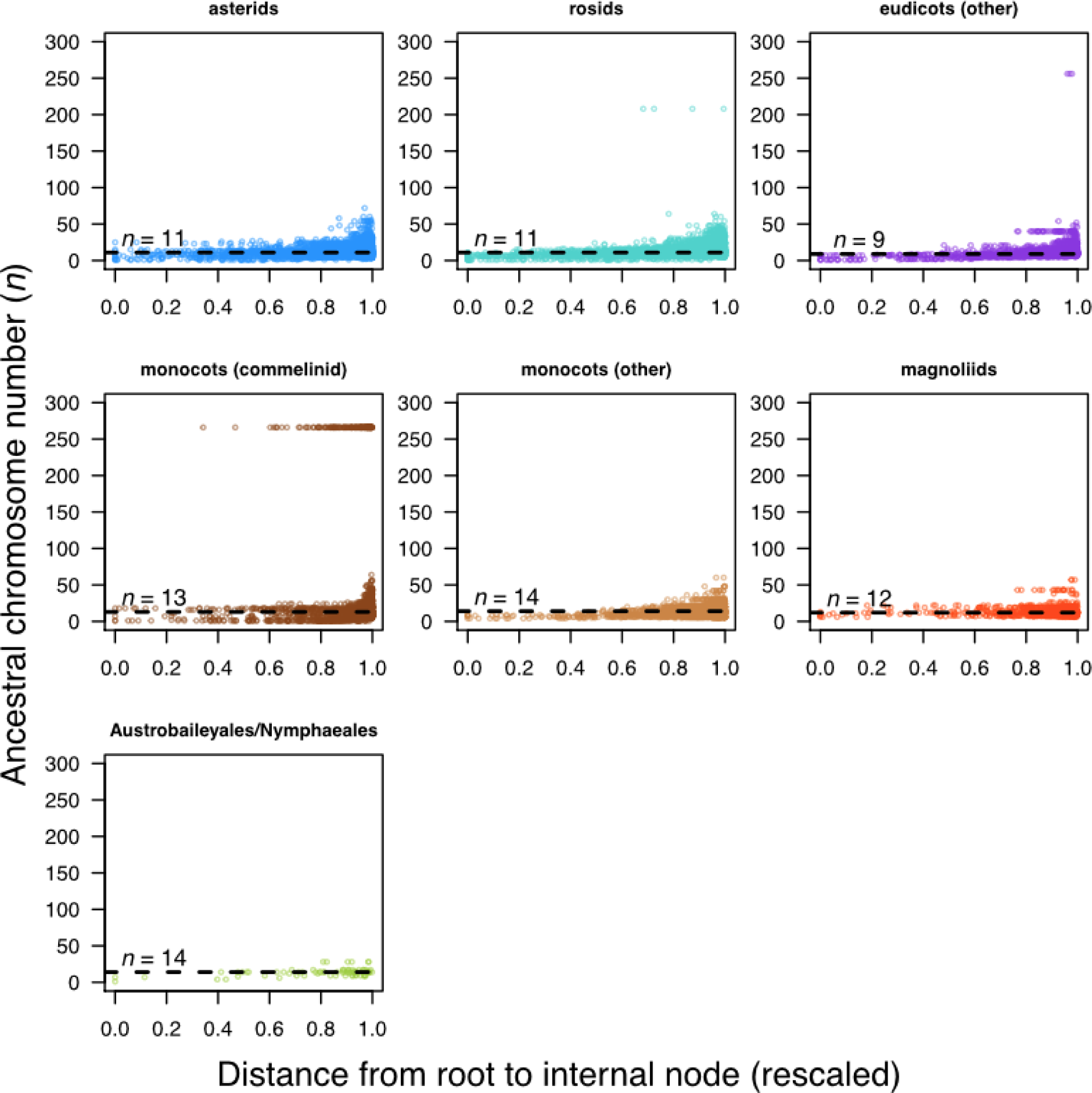
Temporal pattern of chromosome number evolution across major angiosperm groups. The ancestral chromosome numbers were inferred, under the best-fitting ChromEvol model by maximum likelihood, at each internal node in the ***order***-level subtrees extracted from the Qian and Jin (2016) mega-phylogeny. The inferred gametic chromosome numbers at internal nodes are plotted against the age of the node (rescaled such that in each subtree the root depth was one). The dashed horizontal line indicates the median chromosome number across the internal nodes in all the subtrees belonging to a major group.

**Supplementary Figure 3.**
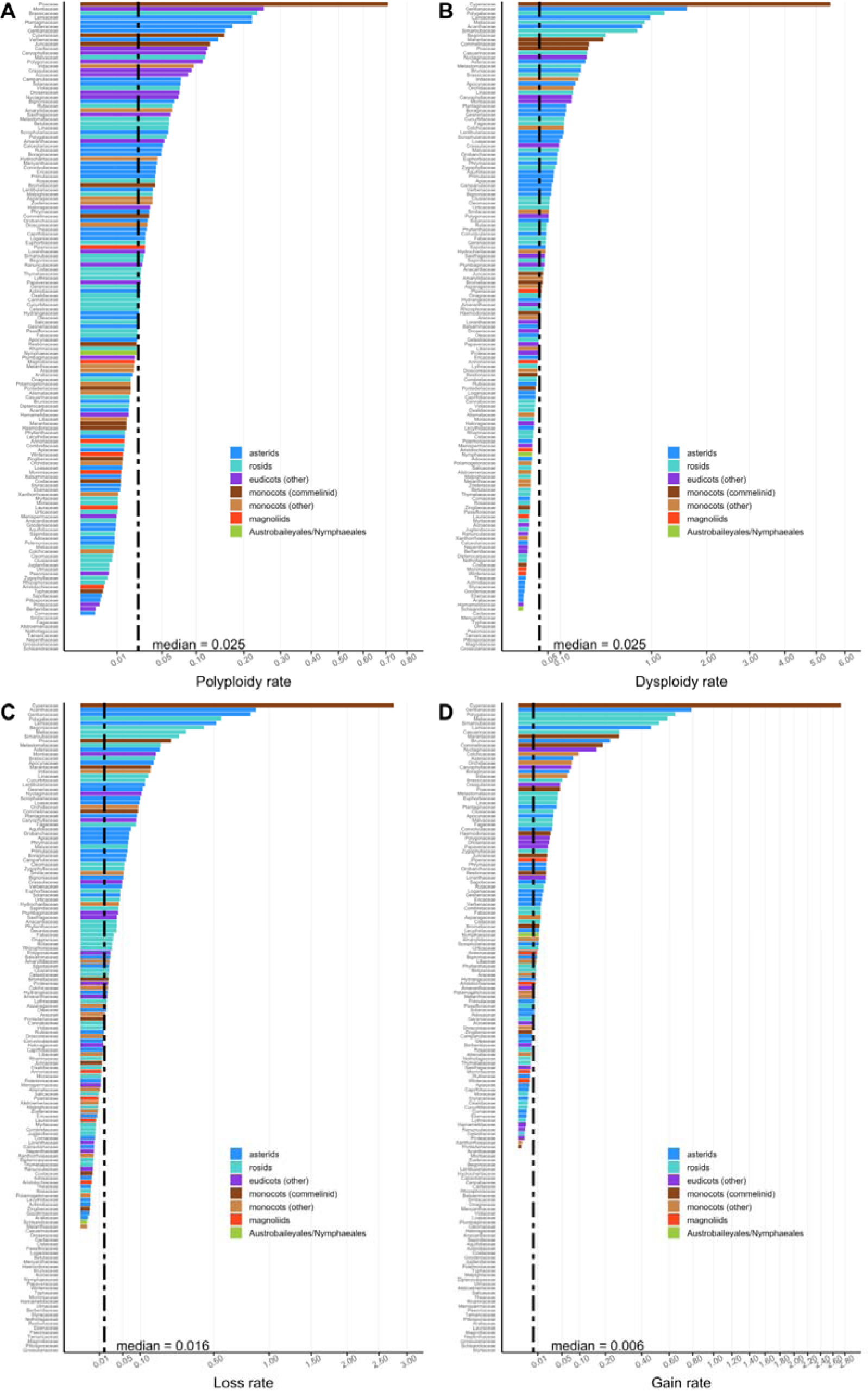
Full version of Figure 2.

**Supplementary Figure 4.**
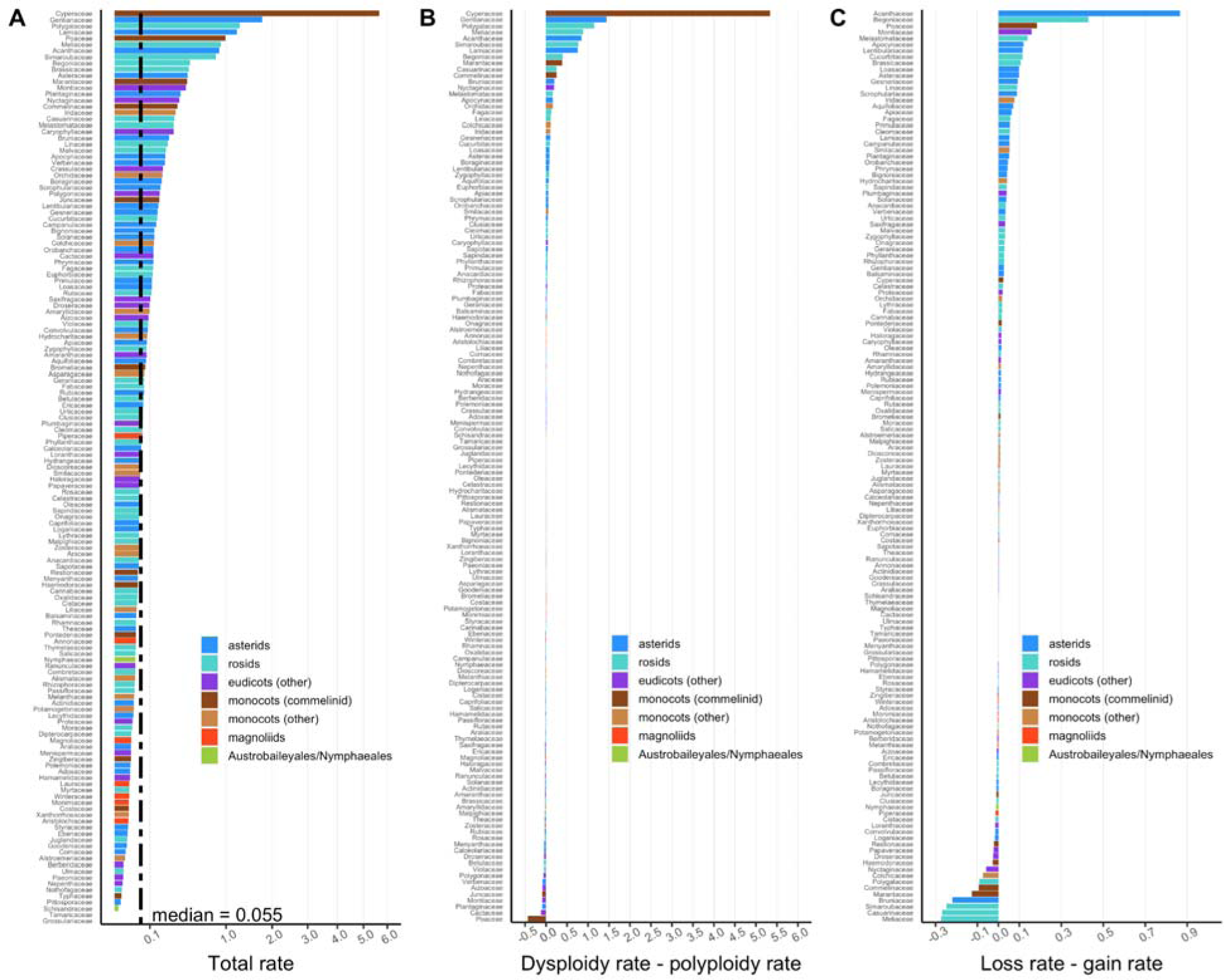
Full version of Figure 4.

**Supplementary Figure 5.**
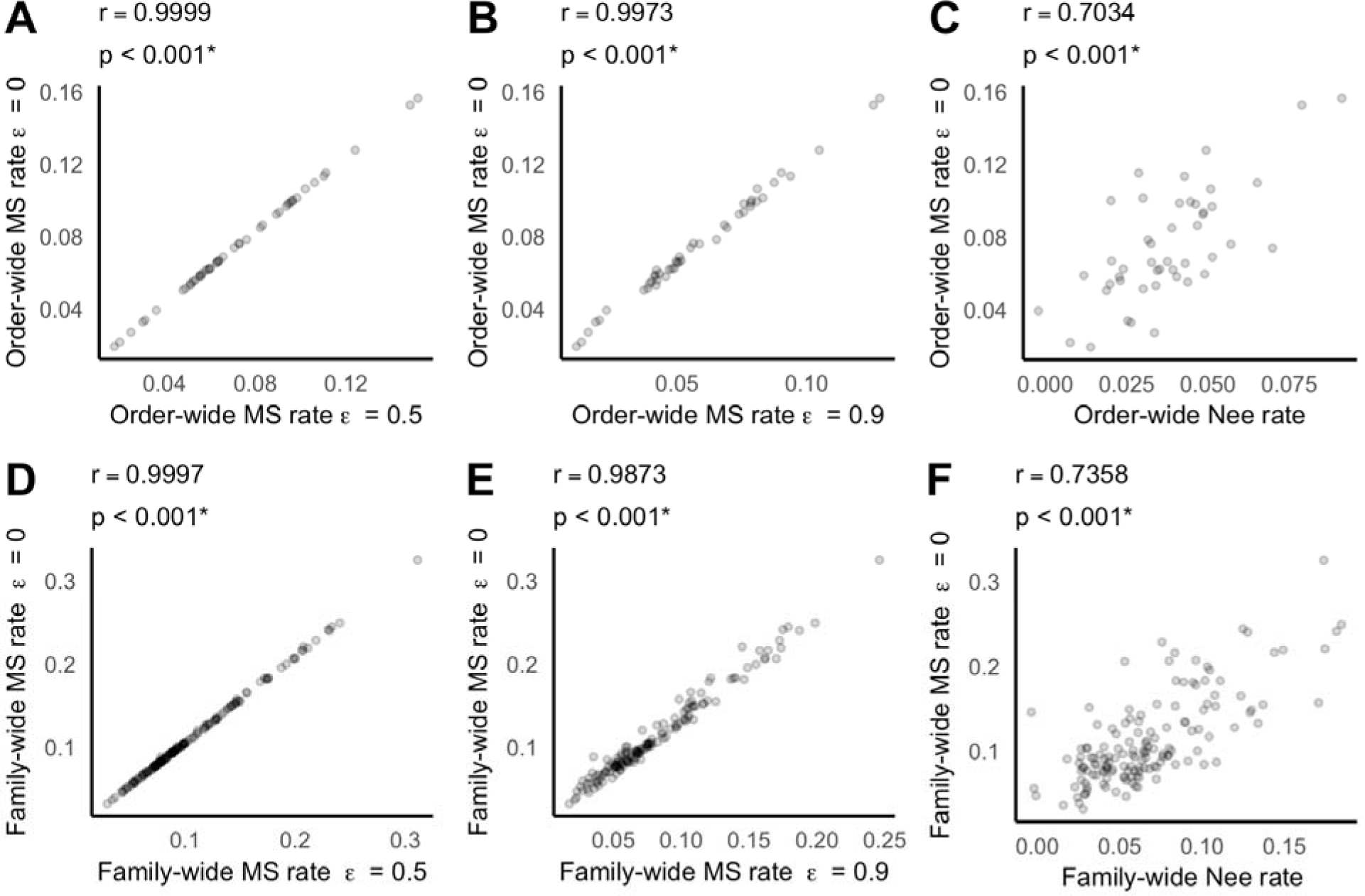
Pairwise correlations between the rates of net diversification estimated using different methods for the angiosperm ***orders*** (A, B, C) and ***families*** (D, E, F). The Magallon and Sanderson (2001) rates were estimated assuming three different values of the relative extinction fraction (ε = 0.00, 0.50, or 0.90). The Nee et al. (1994) rates were estimated under maximum likelihood. Pearson’s correlation coefficients and p-values obtained using the Pearson’s test are shown in the panel titles. An asterisk indicates a p-value of at most 0.05.

**Supplementary Figure 6.**
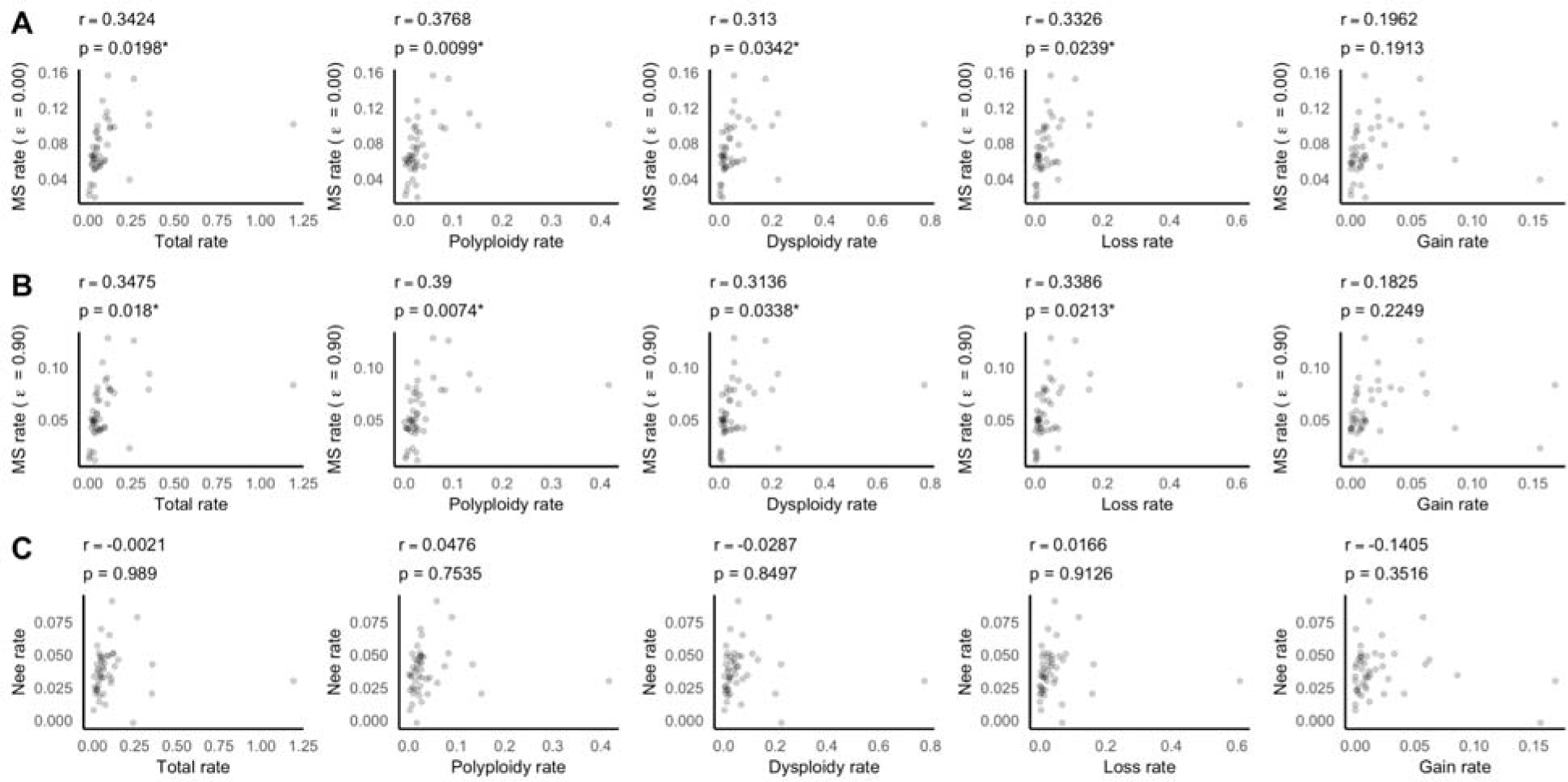
Pairwise correlations between the ***order***-wide rates of net diversification and the rates of chromosome evolution. The Magallon and Sanderson (2001) rates were estimated assuming a relative extinction fraction (ε) of 0.00 (A) or 0.90 (B), and the Nee et al. (1994) rates were estimated under maximum likelihood (C). The rates of chromosomal evolution were estimated under the four-parameter ChromEvol model. The dysploidy rate was computed as the sum of the chromosome gain rate and loss rate. The total rate was computed as the sum of the polyploidy rate and dysploidy rate. Pearson’s correlation coefficients and p-values obtained using the Pearson’s test are shown. An asterisk indicates a p-value of at most 0.05. See **Supplementary Figure 7** for similar plots without Poales (the right most point in each panel).

**Supplementary Figure 7.**
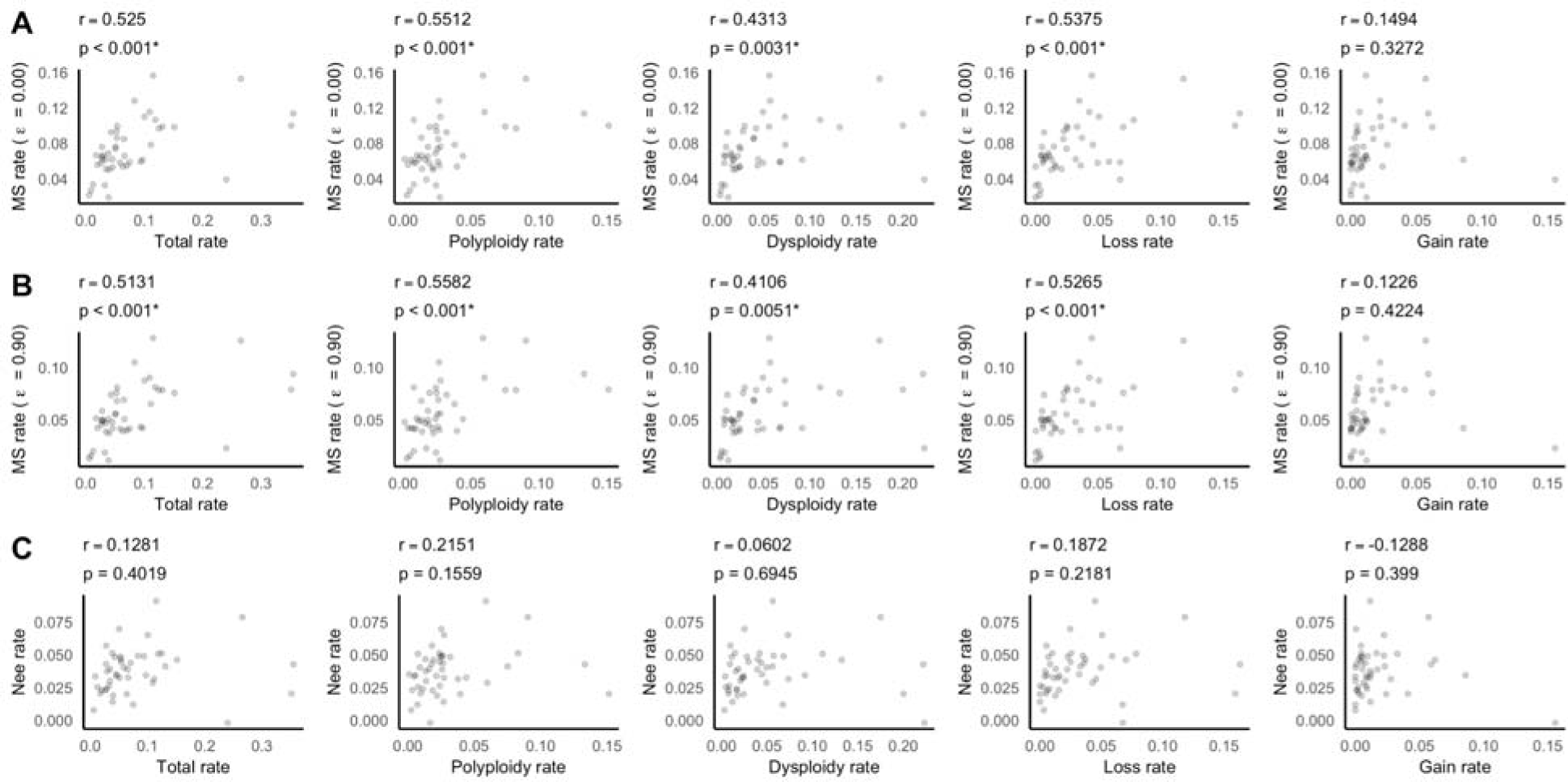
Pairwise correlations between the ***order***-wide rates of net diversification and the rates of chromosomal evolution, *excluding* Poales. See **Supplementary Figure 6** for further explanation.

**Supplementary Figure 8.**
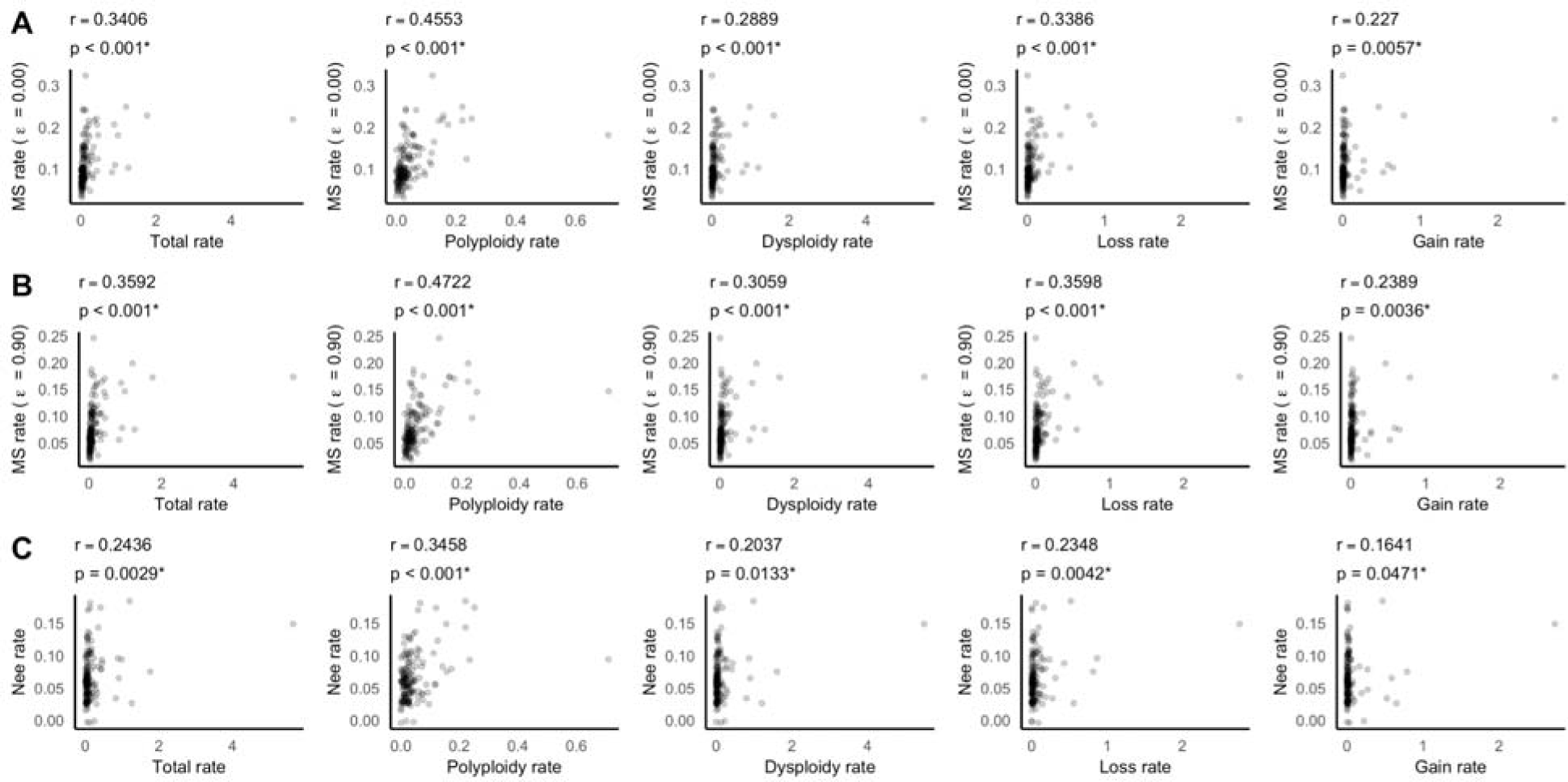
Pairwise correlations between the ***family***-wide rates of net diversification and the rates of chromosomal evolution. See **Supplementary Figure 6** for further explanation and **Supplementary Figure 9** for similar plots without Cyperaceae (the right most points in each panel) and Poaceae, which both have atypically high rates of karyotype evolution.

**Supplementary Figure 9.**
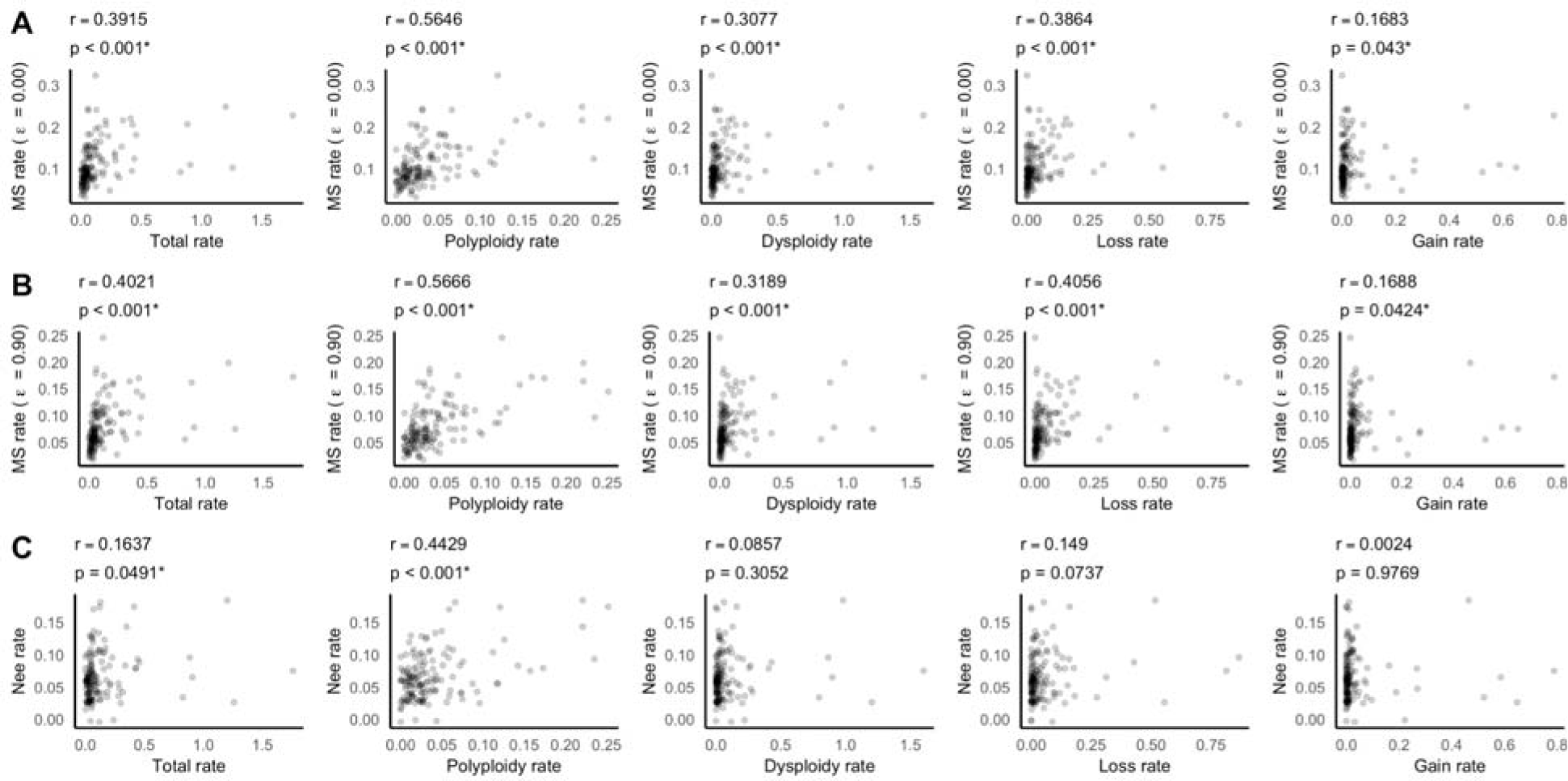
Pairwise correlations between the ***family***-wide rates of net diversification and the rates of chromosomal evolution, *excluding* Cyperaceae and Poaceae. See **Supplementary Figure 6** for further explanation.

**Supplementary Figure 10.**
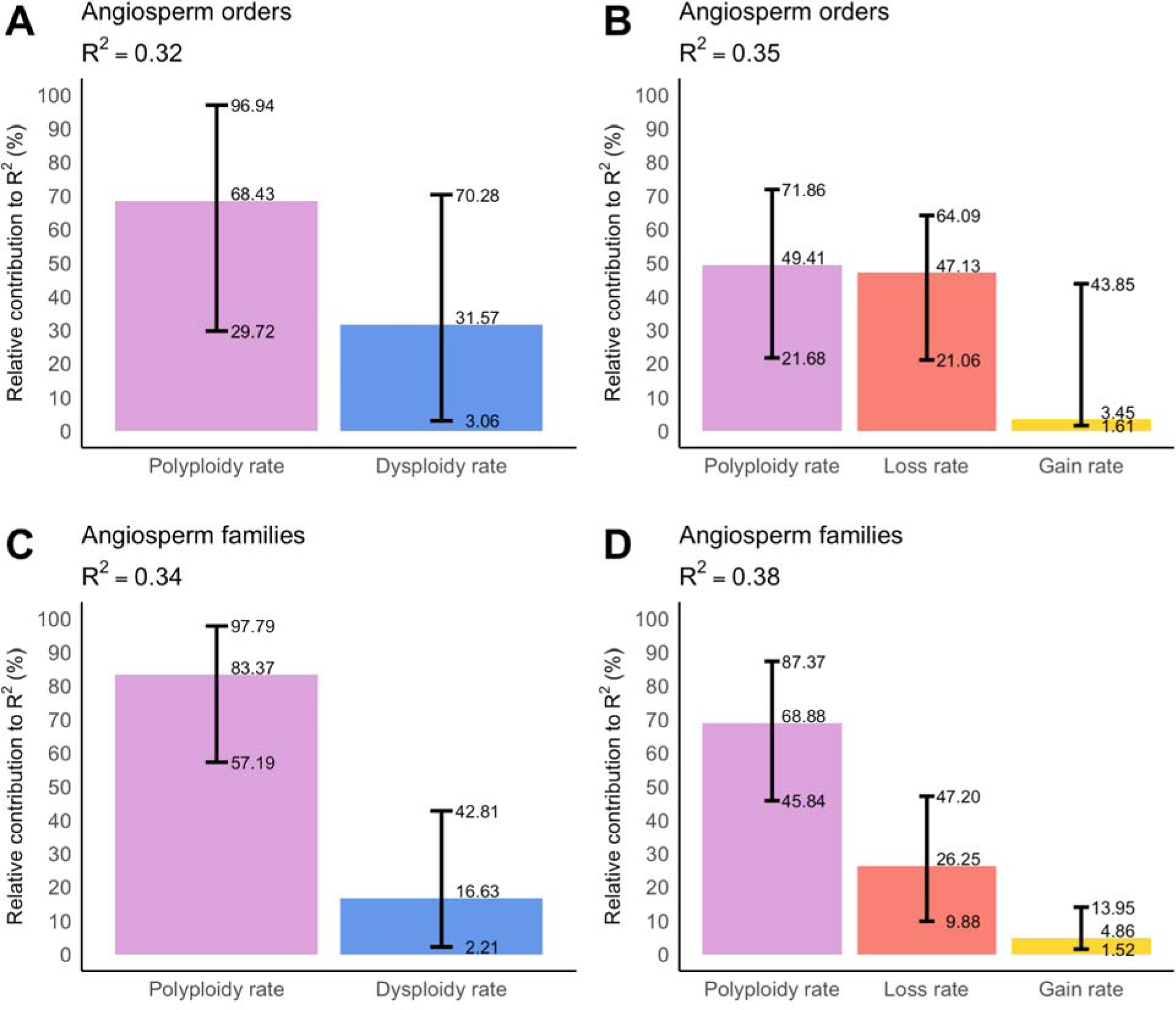
Relative association between the rate of net diversification estimated using the method of Magallon and Sanderson (2001) rate (assuming an extinction rate of zero) and the rates of chromosomal evolution in angiosperm ***orders*** (A, B) and ***families*** (C, D), *excluding* Poales, Cyperaceae, and Poaceae. See Figure 5 for further explanation.

**Supplementary Figure 11.**
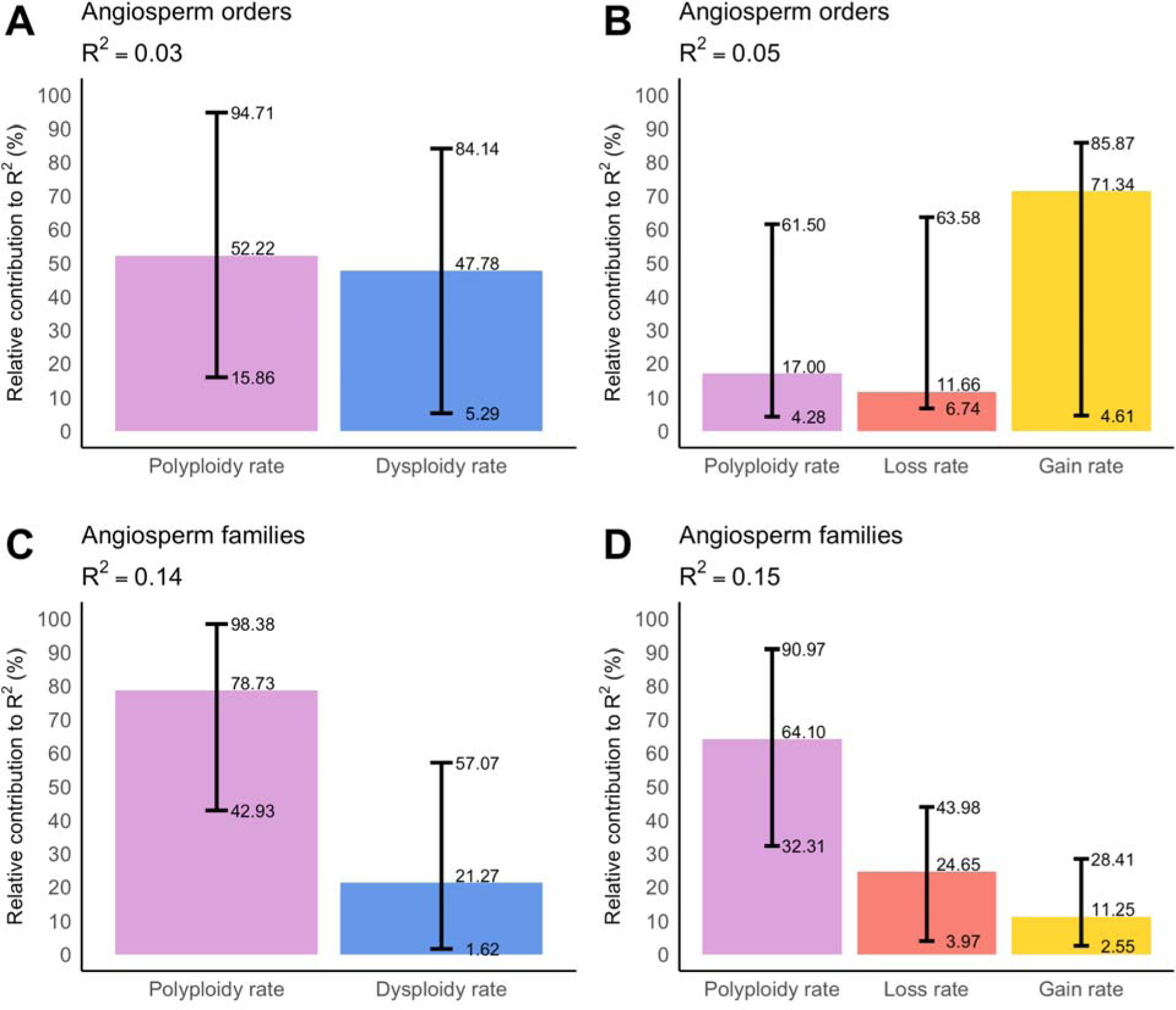
Relative association between the rate of net diversification estimated using the Nee et al. (1994) method and the rates of chromosomal evolution in angiosperm ***orders*** (A, B) and ***families*** (C, D). See Figure 5 for further explanation.

**Supplementary Figure 12.**
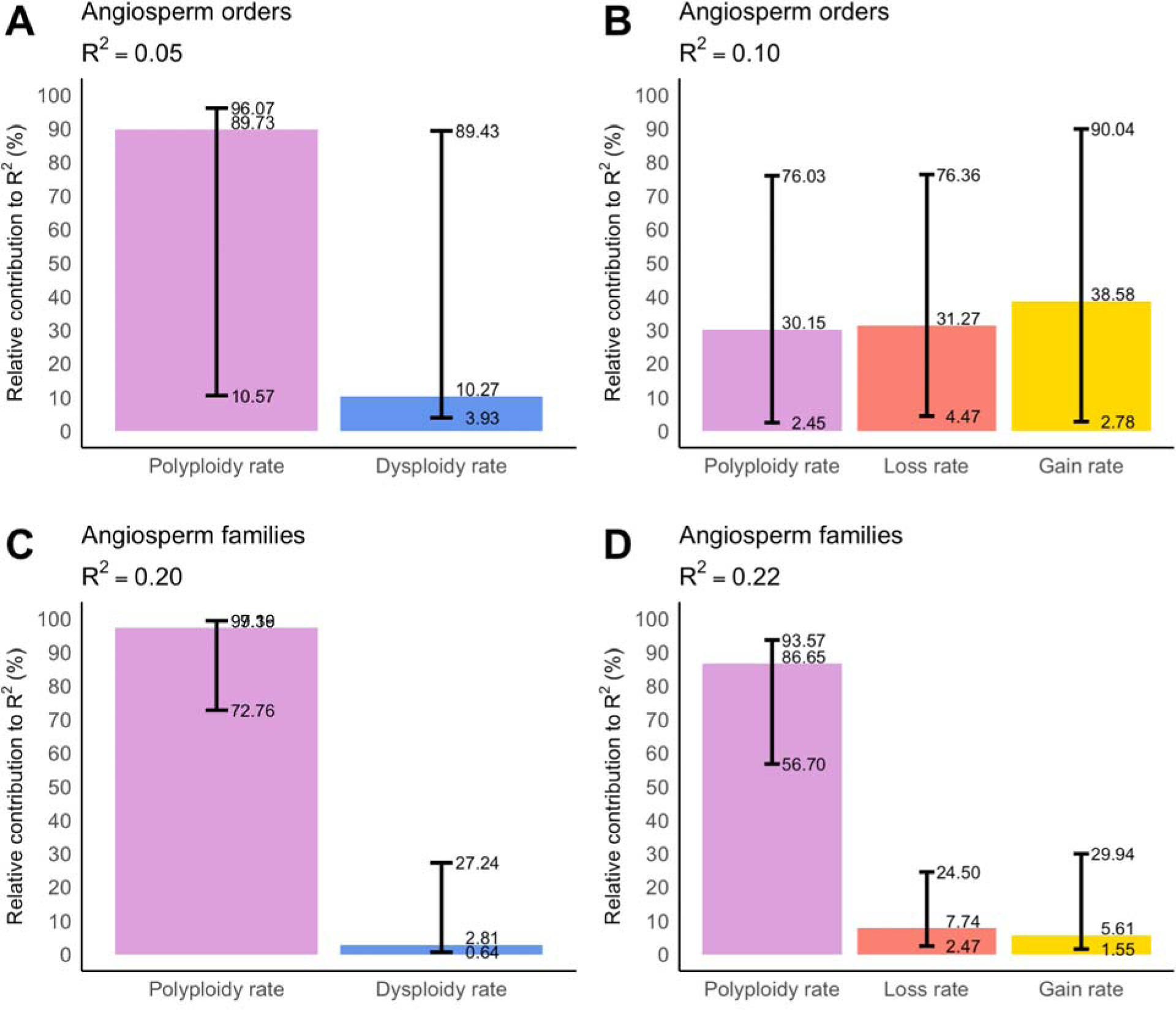
Relative association between the rate of net diversification estimated using the Nee et al. (1994) method and the rates of chromosomal evolution in angiosperm ***orders*** (A, B) and ***families*** (C, D), *excluding* Poales, Cyperaceae, and Poaceae. See Figure 5 for further explanation.

## Notes

### Competing Interest Statement

The authors have declared no competing interest.

## REFERENCES

1. Stebbins, G. L. Chromosomal Evolution in Higher Plants. (Edward Arnold, 1971).

2. Rice, A. et al. The Chromosome Counts Database (CCDB) – a community resource of plant chromosome numbers. New Phytol. 206, 19–26 (2015).

3. Abraham, A. & Ninan, C. A. The chromosomes of Ophioglossum reticulatum L. Curr. Sci. 23, 213–214 (1954).

4. Escudero, M. & Wendel, J. F. The grand sweep of chromosomal evolution in angiosperms. New Phytol. 228, 805–808 (2020).

5. Soltis, P. S. & Soltis, D. E. Ancient WGD events as drivers of key innovations in angiosperms. Curr. Opin. Plant Biol. 30, 159–165 (2016).

6. Otto, S. P. The evolutionary consequences of polyploidy. Cell. 131, 452–462 (2007).

7. Van de Peer, Y., Maere, S. & Meyer, A. The evolutionary significance of ancient genome duplications. Nat. Rev. Genet. 10, 725–732 (2009).

8. te Beest, M. et al. The more the better? The role of polyploidy in facilitating plant invasions. Ann. Bot. 109, 19–45 (2012).

9. Van de Peer, Y., Mizrachi, E. & Marchal, K. The evolutionary significance of polyploidy. Nat. Rev. Genet. 18, 411–424 (2017).

10. Wei, N., Cronn, R., Liston, A. & Ashman, T.-L. Functional trait divergence and trait plasticity confer polyploid advantage in heterogeneous environments. New Phytol. 221, 2286–2297 (2019).

11. Baniaga, A. E., Marx, H. E., Arrigo, N. & Barker, M. S. Polyploid plants have faster rates of multivariate niche differentiation than their diploid relatives. Ecol. Lett. 23, 68–78 (2020).

12. Soltis, D. E. et al. Polyploidy and angiosperm diversification. Am J Bot. 96, 336–348 (2009).

13. Tank, D. C. et al. Nested radiations and the pulse of angiosperm diversification: increased diversification rates often follow whole genome duplications. New Phytol. 207, 454–467 (2015).

14. Landis, J. B. et al. Impact of whole-genome duplication events on diversification rates in angiosperms. Am. J. Bot. 105, 348–363 (2018).

15. One Thousand Plant Transcriptomes Initiative. One thousand plant transcriptomes and the phylogenomics of green plants. Nature 574, 679–685 (2019).

16. Román-Palacios, C., Molina-Henao, Y. F. & Barker, M. S. Polyploids increase overall diversity despite higher turnover than diploids in the Brassicaceae. Proc. Biol. Sci. 287, 20200962 (2020).

17. Mayrose, I. et al. Recently formed polyploid plants diversify at lower rates. Science 333, 1257 (2011).

18. Mayrose, I. et al. Methods for studying polyploid diversification and the dead end hypothesis: a reply to Soltis et al. (2014). New Phytol. 206, 27–35 (2015).

19. Ramsey, J. & Schemske, D. W. Neopolyploidy in flowering plants. Annu. Rev. Ecol. Syst. 33, 589–639 (2002).

20. Fisher, R. A. The sheltering of lethals. Am. Nat. 69, 446–455 (1935).

21. Wright, S. Evolution and the Genetics of Populations, Volume 2: Theory of Gene Frequencies. (University of Chicago Press, 1984).

22. Haldane, J. B. The Causes of Evolution. (Princeton University Press, 1990).

23. Stebbins, G. L. Variation and Evolution in Plants. (Columbia University Press, 1950).

24. Wagner, W. H. Biosystematics and evolutionary noise. Taxon 19, 146–151 (1970).

25. Arrigo, N. & Barker, M. S. Rarely successful polyploids and their legacy in plant genomes. Curr. Opin. Plant Biol. 15, 140–146 (2012).

26. Zenil Ferguson, R. et al. Interaction among ploidy, breeding system and lineage diversification. New Phytol. 224, 1252–1265 (2019).

27. Schubert, I. & Lysak, M. A. Interpretation of karyotype evolution should consider chromosome structural constraints. Trends Genet. 27, 207–216 (2011).

28. Grant, V. Plant Speciation. (Columbia University Press, 1981).

29. Otto, S. P. & Whitton, J. Polyploid incidence and evolution. Annu. Rev. Genet. 34, 401–437 (2000).

30. Márquez-Corro, J. I., Martín-Bravo, S., Spalink, D., Luceño, M. & Escudero, M. Inferring hypothesis-based transitions in clade-specific models of chromosome number evolution in sedges (Cyperaceae). Mol. Phylogenet. Evol. 135, 203–209 (2019).

31. Sader, M. A., Amorim, B. S., Costa, L., Souza, G. & Pedrosa-Harand, A. The role of chromosome changes in the diversification of Passiflora L. (Passifloraceae). Syst. Biodivers. 17, 7–21 (2019).

32. Pimentel, M., Escudero, M., Sahuquillo, E., Minaya, M. Á. & Catalán, P. Are diversification rates and chromosome evolution in the temperate grasses (Pooideae) associated with major environmental changes in the Oligocene-Miocene? PeerJ 5, e3815 (2017).

33. Escudero, M. et al. Karyotypic changes through dysploidy persist longer over evolutionary time than polyploid changes. PLoS ONE 9, e85266 (2014).

34. Zanne, A. E. et al. Three keys to the radiation of angiosperms into freezing environments. Nature 506, 89–92 (2014).

35. Qian, H. & Jin, Y. An updated megaphylogeny of plants, a tool for generating plant phylogenies and an analysis of phylogenetic community structure. *J*. Plant Ecol. 9, 233–239 (2016).

36. Mayrose, I., Barker, M. S. & Otto, S. P. Probabilistic models of chromosome number evolution and the inference of polyploidy. Syst. Biol. 59, 132–144 (2010).

37. Glick, L. & Mayrose, I. ChromEvol: assessing the pattern of chromosome number evolution and the inference of polyploidy along a phylogeny. Mol. Biol. Evol. 31, 1914–1922 (2014).

38. Magallón, S. & Sanderson, M. J. Absolute diversification rates in angiosperm clades. Evolution 55, 1762–1780 (2001).

39. Nee, S., May, R. M. & Harvey, P. H. The reconstructed evolutionary process. Philos. Trans. R. Soc. Lond. B Biol. Sci. 344, 305–311 (1994).

40. Heilborn, O. Chromosome numbers and dimensions, species-formation and phylogeny in the genus Carex. Hereditas 5, 129–216 (2010).

41. Davies, E. W. Cytology, evolution and origin of the aneuploid series in the genus Carex. Hereditas 42, 349–365 (1956).

42. Hipp, A. L., Rothrock, P. E. & Roalson, E. H. The evolution of chromosome arrangements in Carex (Cyperaceae). Bot. Rev. 75, 96–109 (2009).

43. Escudero, M., Hipp, A. L., Waterway, M. J. & Valente, L. M. Diversification rates and chromosome evolution in the most diverse angiosperm genus of the temperate zone (Carex, Cyperaceae). Mol. Phylogenet. Evol. 63, 650–655 (2012).

44. Levy, A. A. & Feldman, M. The impact of polyploidy on grass genome evolution. Plant Physiol. 130, 1587–1593 (2002).

45. Rieseberg, L. H. Chromosomal rearrangements and speciation. Trends in Ecology & Evolution 16, 351–358 (2001).

46. Owens, G. L. & Rieseberg, L. H. Hybrid incompatibility is acquired faster in annual than in perennial species of sunflower and tarweed. Evolution 68, 893–900 (2014).

47. Wood, T. E. et al. The frequency of polyploid speciation in vascular plants. Proc. Natl. Acad. Sci. U. S. A. 106, 13875–13879 (2009).

48. Barker, M. S., Arrigo, N., Baniaga, A. E., Li, Z. & Levin, D. A. On the relative abundance of autopolyploids and allopolyploids. New Phytol. 210, 391–398 (2016).

49. Brochmann, C. et al. Polyploidy in arctic plants. Biol. J. Linn. Soc. 82, 521–536 (2004).

50. Carta, A., Bedini, G. & Peruzzi, L. A deep dive into the ancestral chromosome number and genome size of flowering plants. New Phytol. 228, 1097–1106 (2020).

51. Levin, D. A. & Wilson, A. C. Rates of evolution in seed plants: Net increase in diversity of chromosome numbers and species numbers through time. Proc. Natl. Acad. Sci. U. S. A. 73, 2086–2090 (1976).

52. Zenil-Ferguson, R., Ponciano, J. M. & Burleigh, J. G. Testing the association of phenotypes with polyploidy: An example using herbaceous and woody eudicots. Evolution 71, 1138– 1148 (2017).

53. Stebbins, G. L. Cytological characteristics associated with the different growth habits in the dicotyledons. Am. J. Bot. 25, 189 (1938).

54. Robertson, K., Goldberg, E. E. & Igić, B., Comparative evidence for the correlated evolution of polyploidy and self-compatibility in Solanaceae. Evolution 65, 139–155 (2011).

55. Estep, M. C. et al. Allopolyploidy, diversification, and the Miocene grassland expansion. Proc. Natl. Acad. Sci. U. S. A. 111, 15149–15154 (2014).

56. Thomas, C. D. Rapid acceleration of plant speciation during the Anthropocene. Trends Ecol. Evol. 30, 448–455 (2015).

57. Li, Z. & Barker, M. S. Inferring putative ancient whole-genome duplications in the 1000 Plants (1KP) initiative: access to gene family phylogenies and age distributions. Gigascience 9, (2020).

58. Mandrioli, M. & Manicardi, G. C. Holocentric chromosomes. PLoS Genet. 16, e1008918 (2020).

59. Ruckman, S. N., Jonika, M. M., Casola, C. & Blackmon, H. Chromosome number evolves at equal rates in holocentric and monocentric clades. PLoS Genet. 16, e1009076 (2020).

60. Román-Palacios, C., Medina, C. A., Zhan, S. H. & Barker, M. S. Animal chromosome counts reveal a similar range of chromosome numbers but with less polyploidy in animals compared to flowering plants. Cold Spring Harbor Laboratory 2020.10.10.334722 (2020) doi:10.1101/2020.10.10.334722.

61. Wendel, J. F. The wondrous cycles of polyploidy in plants. Am. J. Bot. 102, 1753–1756 (2015).

62. Dodsworth, S., Chase, M. W. & Leitch, A. R. Is post-polyploidization diploidization the key to the evolutionary success of angiosperms? Bot. J. Linn. Soc. 180, 1–5 (2015).

63. Mandáková, T., Li, Z., Barker, M. S. & Lysak, M. A. Diverse genome organization following 13 independent mesopolyploid events in Brassicaceae contrasts with convergent patterns of gene retention. Plant J. 91, 3–21 (2017).

64. Mandáková, T. et al. Multispeed genome diploidization and diversification after an ancient allopolyploidization. Mol. Ecol. 26, 6445–6462 (2017).

65. Mandáková, T. & Lysak, M. A. Post-polyploid diploidization and diversification through dysploid changes. Curr. Opin. Plant Biol. 42, 55–65 (2018).

66. Li, Z. et al. Patterns and processes of diploidization in land plants. Annu. Rev. Plant Biol. (2021) doi:10.1146/annurev-arplant-050718-100344.

67. Fawcett, J. A., Maere, S. & Van de Peer, Y. Plants with double genomes might have had a better chance to survive the Cretaceous-Tertiary extinction event. Proc. Natl. Acad. Sci. U. S. A. 106, 5737–5742 (2009).

68. Chen, Z. J. Molecular mechanisms of polyploidy and hybrid vigor. Trends Plant Sci. 15, 57–71 (2010).

69. Dijkstra, H. & Speckmann, G. J. Autotetraploidy in caraway (Carum carvi L.) for the increase of the aetheric oil content of the seed. Euphytica 29, 89–96 (1980).

70. Lavania, U. C. & Srivastava, S. Enhanced productivity of tropane alkaloids and fertility in artificial autotetraploids of *Hyoscyamus niger* L. Euphytica 52, 73–77 (1991).

71. Hull Sanders, H. M. & Johnson, R. H. Effects of polyploidy on secondary chemistry, physiology, and performance of native and invasive genotypes of Solidago gigantea (Asteraceae). Am. J. Bot. 96, 762–770. (2009).

72. Li, W.-L., Berlyn, G. P. & Ashton, P. M. S. Polyploids and their structural and physiological characteristics relative to water deficit in Betula papyrifera (Betulaceae). Am. J. Bot. 83, 15–20 (1996).

73. Maherali, H., Walden, A. E. & Husband, B. C. Genome duplication and the evolution of physiological responses to water stress. New Phyt. 184, 721–731 (2009).

74. Fawcett, J. A. & Van de Peer, Y. Angiosperm polyploids and their road to evolutionary success. Trends Evol. Biol. 1, e3 (2010).

75. Novikova, P. Y. et al. Polyploidy breaks speciation barriers in Australian burrowing frogs *Neobatrachus*. PLoS Genet. 16, e1008769 (2020).

76. Scarpino, S. V., Levin, D. A. & Meyers, L. A. Polyploid formation shapes flowering plant diversity. Am. Nat. 184, 456–465 (2014).

77. Meudt, H. M. et al. Is genome downsizing associated with diversification in polyploid lineages ofVeronica? Bot. J. Linn. Soc. 178, 243–266 (2015).

78. Schranz, M. E., Eric Schranz, M., Mohammadin, S. & Edger, P. P. Ancient whole genome duplications, novelty and diversification: the WGD Radiation Lag-Time Model. Curr. Opin. Plant Biol. 15, 147–153 (2012).

79. Márquez-Corro, J. I., Escudero, M. & Luceño, M. Do holocentric chromosomes represent an evolutionary advantage? A study of paired analyses of diversification rates of lineages with holocentric chromosomes and their monocentric closest relatives. Chromosome Res. 26, 139–152 (2018).

80. Freyman, W. A. & Höhna, S. Cladogenetic and anagenetic models of chromosome number evolution: a Bayesian model averaging approach. Syst. Biol. 67, 195–215 (2018).

81. Zenil-Ferguson, R. et al. Interaction among ploidy, breeding system and lineage diversification. New Phytol. 224, 1252–1265 (2019).

82. Paradis, E., Claude, J. & Strimmer, K. APE: Analyses of Phylogenetics and Evolution in R language. Bioinformatics 20, 289–290 (2004).

83. The Plant List. The Plant List http://www.theplantlist.org/.

84. Kluyver, T. A. & Osborne, C. P. Taxonome: a software package for linking biological species data. Ecol. Evol. 3, 1262–1265 (2013).

85. Angiosperm Phylogeny Website. http://www.mobot.org/MOBOT/research/APweb/.

86. Akaike, H. A new look at the statistical model identification. IEEE Trans. Automat. Contr. 19, 716–723 (1974).

87. Scholl, J. P. & Wiens, J. J. Diversification rates and species richness across the Tree of Life. Proc. Biol. Sci. 283, (2016).

88. Pennell, M. W. et al. geiger v2.0: an expanded suite of methods for fitting macroevolutionary models to phylogenetic trees. Bioinformatics 30, 2216–2218 (2014).

89. FitzJohn, R. G. Diversitree : comparative phylogenetic analyses of diversification in R. Methods Ecol. Evol. 3, 1084–1092 (2012).

90. Schliep, K. P. phangorn: phylogenetic analysis in R. Bioinformatics 27, 592–593 (2011).

91. Revell, L. J. phytools: an R package for phylogenetic comparative biology (and other things). Methods Ecol. Evol. 3, 217–223 (2012).

92. Rowan, T. H. Functional stability analysis of numerical algorithms. (1991).

93. King, A. A. *subplex*. (Github).

94. Huber, P. J. Robust Statistics. (John Wiley & Sons, Inc., 1981).

95. Venables, W. N. & Ripley, B. D. Modern Applied Statistics with S. (Springer-Verlag New York, 1994).

96. Lindeman, R. H., Merenda, P. F. & Gold, R. Z. Introduction to Bivariate and Multivariate Analysis. (Scott, Foresman, 1980).

97. Grömping, U. Relative importance for linear regression in R: the package *relaimpo*. J. Stat. Soft. 17 (2006).

98. Grömping, U. Estimators of relative importance in linear regression based on variance decomposition. Am. Stat. 61, 139–147 (2007).

